# Targeting a broad spectrum of *KRAS*-mutant cancers by hyperactivation-induced cell death

**DOI:** 10.1101/2022.09.21.508660

**Authors:** Johanna Lilja, Jasmin Kaivola, James R.W. Conway, Joni Vuorio, Hanna Parkkola, Pekka Roivas, Taru Varila, Guillaume Jacquemet, Emilia Peuhu, Emily Wang, Ulla Pentikäinen, Itziar Martinez D. Posada, Hellyeh Hamidi, Arafat K. Najumudeen, Owen J. Sansom, Igor L. Barsukov, Daniel Abankwa, Ilpo Vattulainen, Marko Salmi, Johanna Ivaska

**Affiliations:** Turku Bioscience Centre, University of Turku, FI-20520 Turku, Finland; Department of Physics, University of Helsinki, Helsinki, Finland; Institute of Biomedicine, University of Turku, FI-20520 Turku, Finland; Faculty of Science and Engineering, Cell Biology, Åbo Akademi University, FI-20520 Turku, Finland; Institute of Biomedicine, Cancer Research Laboratory FICAN West, University of Turku, FIN-20520 Turku, Finland; University of Liverpool, Liverpool, United Kingdom; CRUK Beatson Institute, Garscube Estate, Switchback Road, Glasgow, G61 1BD, United Kingdom; Institute of Cancer Sciences, University of Glasgow, Garscube Estate, Switchback Road, Glasgow, G61 1QH, United Kingdom; Department of Life Sciences and Medicine, University of Luxembourg, 4365 Esch- sur-Alzette, Luxembourg; MediCity Research Laboratory, University of Turku, FI-20520 Turku, Finland; Department of Life Technologies, University of Turku, Turku, Finland; InFLAMES Research Flagship Center, University of Turku, Turku, Finland.; Foundation for the Finnish Cancer Institute, Tukholmankatu 8, FI-00014 Helsinki

## Abstract

The *KRAS* oncogene drives many common and highly fatal malignancies. These include pancreatic, lung, and colorectal cancer, where numerous different activating *KRAS* mutations have made the development of KRAS inhibitors difficult. Here we identify the scaffold protein SH3 and multiple ankyrin repeat domain 3 (SHANK3) as a RAS interactor that binds overactive mutant forms to limit oncogenic KRAS signalling and maintain RAS-activity at an optimal level. Depletion of SHANK3 results in hyperactivation of KRAS/mitogen-activated protein kinase (MAPK) signalling, which in turn selectively induces MAPK/ERK-dependent cell death in *KRAS*-mutant cancers. Furthermore, targeting of this therapeutic vulnerability through nanobody- or RNA interference-mediated disruption of the SHANK3-KRAS interaction reduces tumour growth *in vivo.* Thus, inhibition of the SHANK3-KRAS interaction represents a new pan-KRAS-mutant compatible strategy for selective killing of *KRAS*- mutant cancer cells through excessive signalling.

**Graphical abstract:** 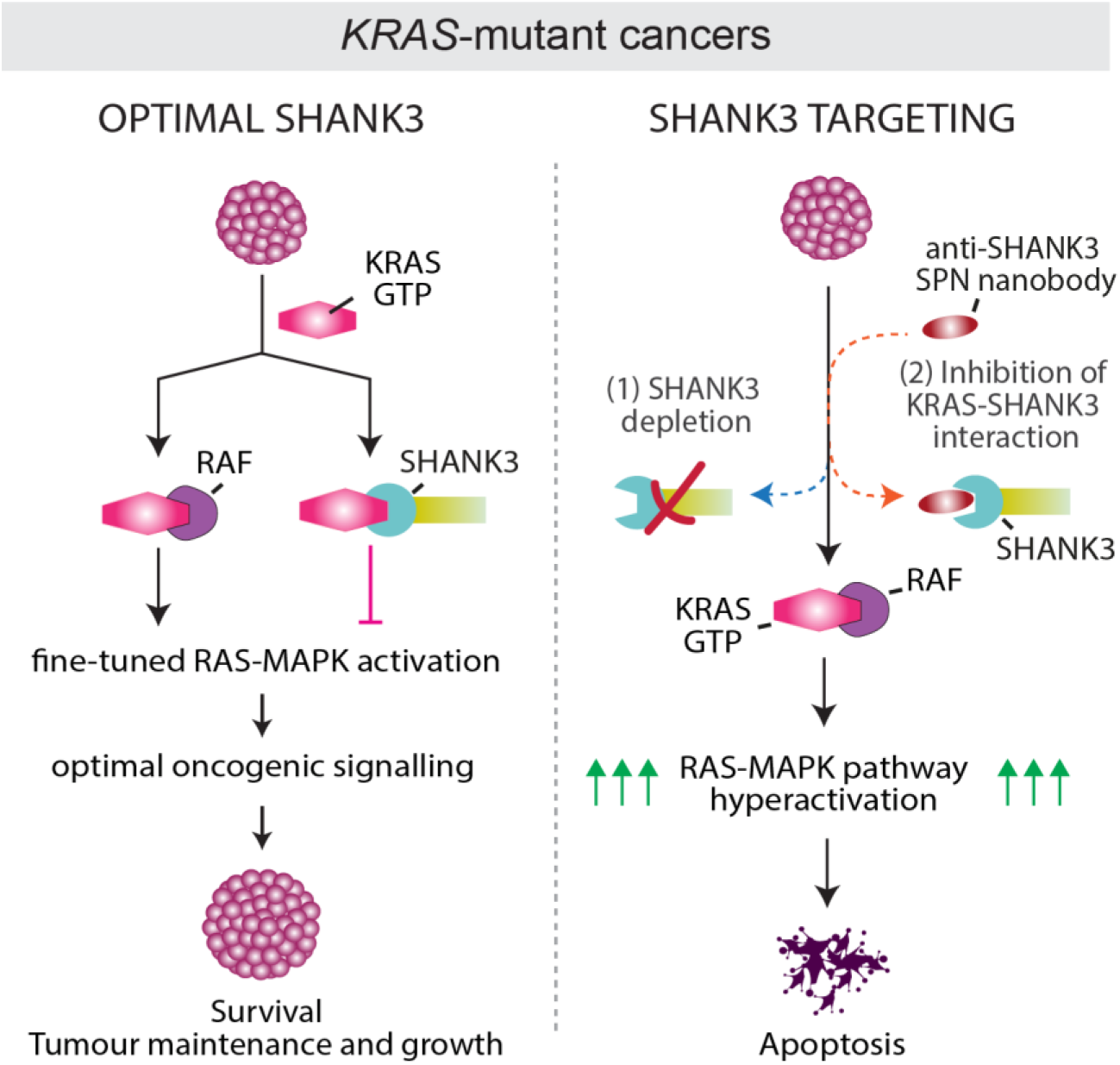

Schematic model of SHANK3-controlled cell fate in *KRAS*-mutant cancers. SHANK3 directly interacts with KRAS and competes with RAF for KRAS binding to sustain oncogenic RAS-MAPK/ERK signalling at an optimal level (i.e. below toxic oncogenic signalling) in *KRAS*-mutant cancers. SHANK3 silencing (1) and inhibition of SHANK3-KRAS interaction (2) drive *KRAS*-mutant cells into cell death.

## Introduction

Aberrant activity of KRAS has been identified in more than 20% of all human cancers (*1*). The incidence is substantially higher in some of the most inherently therapy-resistant cancers, including non-small cell lung cancers (NSCLC; 30% incidence), colorectal cancers (CRC; 50% incidence), and pancreatic ductal adenocarcinomas (PDAC; 95% incidence) (*1, 2*). Oncogenic mutations in *KRAS* induce the constitutive activation of proliferative signalling cascades, promoting cancer progression and also conferring resistance to standard first-line treatments (*3–5*). Unfortunately, survival of patients with *KRAS*-mutated NSCLC and PDAC has barely improved over the past few decades (*6*), highlighting an urgent need to broaden our view on how to target oncogenic KRAS.

KRAS is a plasma membrane associated small GTPase, active in its GTP-bound state and inactive in the GDP-bound state (*7*). Active KRAS interacts with downstream effectors, such as RAF, which in turn activate, for example, the MAPK/extracellular-signal-regulated kinase (ERK) pathway leading to apoptosis inhibition and activation of transcription factors that promote cancer cell survival and growth (*8, 9*). Oncogenic forms of KRAS are predominantly in the GTP-bound active state, supporting high-level effector pathway activation to drive excessive cell proliferation and survival (*10*).

KRAS mutations have a long history of resistance to drug therapy. However, through the accrued knowledge of the structural and biochemical characteristics of these mutants, the field has seen a recent development of mutation-specific drugs with promising preclinical and clinical efficacy (*10–18*). The most promising mutation-specific inhibitors are those targeting KRASG12C, a mutation found in ∼12% of all KRAS-driven tumours (*11–16, 19, 20*). Therefore, the clinical promise of these inhibitors only applies to the subset of patients carrying the given mutation, leaving ∼88% of patients without a KRAS-targeted treatment option. In addition, various resistance mechanisms have already been reported, indicating significant limitations of mutation-specific inhibitors in heterogenous tumours (*21–23*).

To address a broader patient population, a vast effort is being made in several new pan-KRAS approaches targeting all KRAS mutants (*10, 24*). These include pharmacological manipulation of KRAS upstream activators, such as SHP2 and SOS1 (*25–27*) or development of pan-KRAS protein degradation strategies, such as proteolysis-targeting chimeras (PROTACs) (*28*). Pan-KRAS inhibitors hold promise for patients for whom targeted therapy remains elusive. Therefore, there is an unmet need to identify new vulnerabilities associated with KRAS-dependent tumours and introduce novel approaches that broadly target these cancers, regardless of the activating mutation.

We and others previously showed that the multidomain scaffold protein SH3 and multiple ankyrin repeat domain 3 (SHANK3), adopts a RAS-association (RA) domain-like fold with high affinity for GTP-bound RAS and Rap G-proteins with its N-terminal Shank/ProSAP (SPN) –domain (*29, 30*). SHANK3 was initially identified in excitatory synapses of the central nervous system (*31*), but we have demonstrated a role for SHANK3 in regulation of the cytoskeleton (*32*) and in cell adhesion through binding Rap1 and inhibiting integrins in non-neuronal cells and cancer cells (*29*). However, whether SHANK3 plays a functional role in oncogenic RAS signalling remains unknown.

In the present study, our aim was to elucidate whether SHANK3 plays a functional role in KRAS-driven cancers. Using several different approaches, we identify SHANK3 as a RAS interactor that competes with RAF for binding to active mutant forms of KRAS and subsequently limits oncogenic RAS-MAPK signalling. We show that SHANK3 depletion results in hyperactivation of KRAS signalling and induces cell death in a broad range of *KRAS*-mutant cancers *in vitro* and *in vivo*. Our data demonstrate that SHANK3 depletion in pre-existing *KRAS*- mutant tumours impairs tumourigenic growth, highlighting the potential of SHANK3 targeting as an anti-cancer therapy. To provide proof-of-concept evidence that SHANK3 can be targeted pharmacologically, we developed nanobodies disrupting the SHANK3-KRAS protein-protein interaction and demonstrate their efficacy in inducing apoptosis in *KRAS*-mutant cancers. Collectively our data reveal that the SHANK3-KRAS interaction is an exploitable vulnerability of pan-KRAS-driven cancer.

## Results

### *SHANK3* depletion impairs cell proliferation in a large panel of *KRAS*-mutant cancer cell lines

The Cancer Dependency Map and our current view of genes essential for cancer survival are largely based on the pan-cancer Broad and Sanger CRISPR-Cas9 viability screens and genome-scale RNA interference screens such as Achilles, which did not include guide RNAs or shRNA against *SHANK3* (*33, 34*). Therefore, to investigate the role of SHANK3 in cancer cell viability we depleted endogenous *SHANK3* using two unique RNAi oligonucleotides in a large panel of human PDAC, NSCLC and CRC cell lines harbouring either distinct *KRAS* mutations or wild-type (WT) *KRAS* (Fig. 1a and silencing validated in Extended Data Fig. 1a). Cell proliferation was strongly impaired with both *SHANK3* siRNAs in each of the 12 tested cancer cell lines with activating mutations in *KRAS* [mean inhibition (%): 63.3±4.4% (PANC-1), 66.1±5.6% (Panc10.05), 38.9±4.2% (AsPC-1), 63.0±19.6% (Su86.86), 55.5±4.7% (SW1990), 55.5±8.8% (YAPC), 74.5±4.6% (PaTu8902), 79.5±19.1% (MIA PaCa-2), 50.8±20.5% (A549), 72.1±7.5% (H441), 38.0±22.1% (HCT-15), 81.3±8.6% (HCT-116)] (Fig. 1a). In contrast, the tested cancer cell lines harboring WT *KRAS* did not show significant co-directional inhibition by the two unique SHANK3 siRNAs [mean inhibition (%): -3.9±33.2% (H292), 11.8±12.3% (H266), 14.0±20.9% (H226), 13.6±8.0% (HT-29)] (Fig. 1a). Non-transformed epithelial cells showed no response to SHANK3 inhibition [mean inhibition (%): 0.4±2.5%]. These data indicate low or no sensitivity to *SHANK3* depletion in non-*KRAS* mutated cells (Fig. 1a).

**Figure 1.**
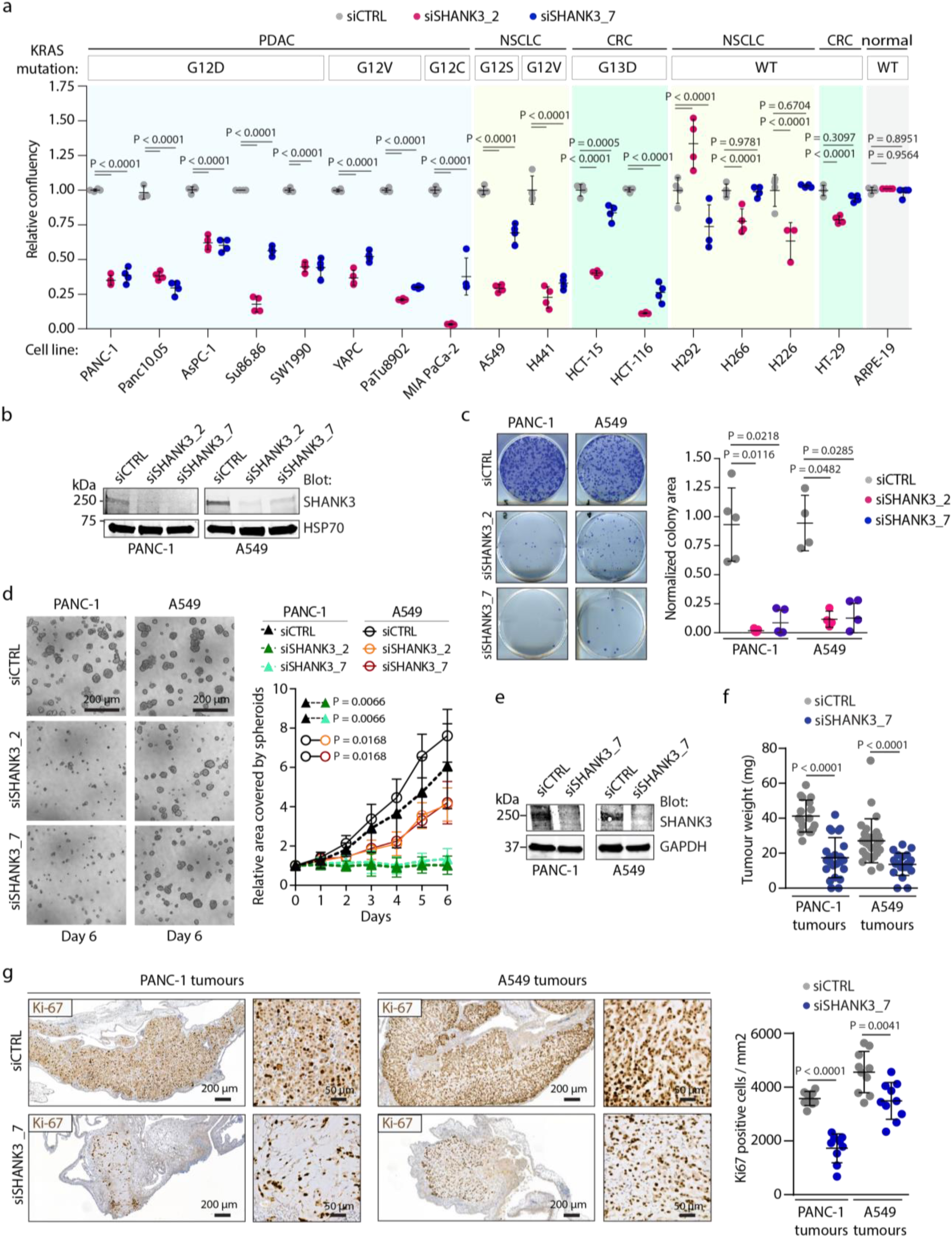
SHANK3 depletion inhibits cell proliferation *in vitro* and *in vivo* in different cancer types driven by distinct *KRAS* mutations. a, Cell proliferation screen in pancreatic (PDAC), lung (NSCLC) and colorectal (CRC) cancer cell lines with the indicated *KRAS* mutations after control (siCTRL, grey) and *SHANK3*-silencing (siSHANK3_2 (red) or siSHANK3_7 (blue)). Cancer lines with wild-type (WT) *KRAS* and non-transformed cells (normal) were also analysed. Shown are the individual data points and the mean ± s.d. (the mean of the control is set to 1.0 by definition; n = 3-4). Dunnett’s multiple comparisons test. b, Immunoblotting of SHANK3 protein levels in the siRNA-silenced *KRAS*-mutant PANC-1 (KRASG12D) and A549 (KRASG12S) cells. HSP70 is a loading control. c, Colony growth of control or *SHANK3*-silenced PANC-1 and A549 cells. Shown are representative images and quantification of the colony areas (individual data points and the mean ± s.d.); n = 5 (PANC-1) and 4 (A549) independent experiments. Kruskal-Wallis test and Dunn’s post hoc test. d, Spheroid growth of control or *SHANK3*-silenced PANC-1 or A549 cells. Shown are representative images and quantification (mean ± s.d) of the spheroid areas; n = 3 independent experiments. One-way ANOVA with Holm-Sidak’s multiple comparison test (at the endpoint). e-g, Tumour growth of control or *SHANK3*-silenced PANC-1 and A549 cells on CAM membranes. e, Immunoblotting of SHANK3 protein levels. f, Tumour weights. g, Representative immunostainings and quantification of Ki-67 positive cells in tumours. Shown are individual data points and mean ± s.d. Data are from 17-28 (f) and 9-10 (g) tumours per sample group. Mann Whitney test (f) and Unpaired Student’s t-test with Welch’s correction (g).

Acute *SHANK3* silencing with two independent siRNAs also reduced colony growth of *KRAS*-mutant pancreatic (PANC-1) and lung cancer (A549) cell lines by approximately 90% (Fig. 1b,c). Similarly, the growth of *KRAS*-mutant pancreatic (PANC-1 and AsPC-1) and lung cancer (A549) cells was significantly reduced in a 3- dimensional (3D) cancer spheroid model (Fig. 1d and Extended Data Fig. 1b, c).

To examine if SHANK3 is also essential for tumourigenesis *in vivo*, we established PDAC and LUAD xenograft models on chick embryo chorioallantoic membrane (CAM) in fertilised eggs. In line with our *in vitro* results, we observed a significant decrease both in tumour weight and in the number of proliferating Ki-67+ cells in *SHANK3*-silenced KRAS-driven PANC-1 and A549 xenografts when compared to control-silenced tumours (Fig. 1e-g and Extended Data Fig. 1d).

These data collectively indicate that the depletion of endogenous SHANK3 effectively blocks cell proliferation and growth *in vitro* and *in vivo* in different cancer types driven by distinct *KRAS* mutations.

### SHANK3 interacts specifically with active KRAS to regulate *KRAS*-mutant cell survival

To investigate the molecular mechanisms of SHANK3-dependent suppression of KRAS-driven tumour growth, we first analysed whether this could be linked to SHANK3 interaction with active/mutant KRAS. We previously determined the crystal structure of the N-terminal part of the SHANK3 protein [comprising of SPN and ARR (ankyrin repeat region) domains; Fig. 2a] revealing that the SPN-domain adopts a RAS-association (RA) domain-like structure (29). Microscale thermophoresis (MST) and isothermal titration calorimetry (ITC) measurements with purified recombinant proteins indicated that the N-terminal SHANK3 SPN-domain alone and the SPN-ARR fragment (Fig. 2a) interacted with active (GMPPCP-form) KRAS-mutants with similar affinities (Kd = 5.0 ± 0.6 μM for G12V with SPN; and Kd = 5.4 ± 0.7 μM for Q61H with SPN-ARR; Fig. 2b,c and Extended Data Fig. 2a-d). Inactive (GDP-bound) KRAS showed no interaction with SHANK3 SPN-ARR in ITC measurements (Fig. 2d), indicating that the interactions were specific to GTP-bound KRAS. These data show that the SPN domain of SHANK3 directly interacts with KRAS in an activity dependent manner similar to established RA domain-containing proteins.

**Figure 2.**
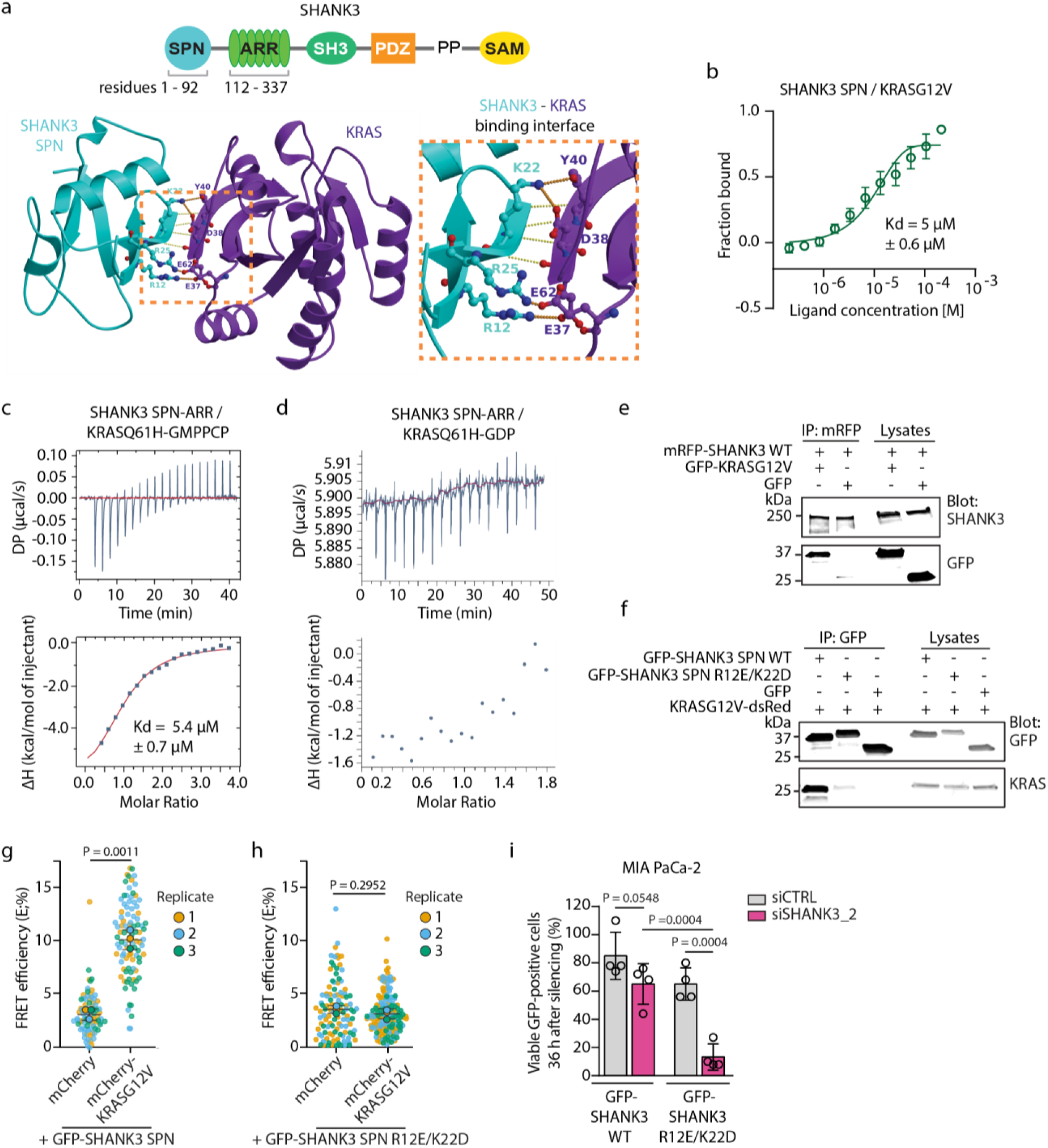
SHANK3 directly interacts with active KRAS, regardless of the activating mutation. **a,** Top: Schematic model of SHANK3 protein domains. SPN, Shank/ProSAP N-terminal domain; ARR, ankyrin repeat domain; SH3, Src homology 3 domain; PDZ, PSD-95/Discs large/ZO-1 domain; PP, proline-rich region; SAM, sterile alpha motif domain. Bottom: Crystal-structure based model of SHANK3 SPN domain in complex with active KRAS. The zoom-in shows the critical interacting amino acids. The SHANK3 SPN-KRAS interaction is disrupted by R12E/K22D mutation in SHANK3. b, Microscale thermophoresis (MST) binding curve for the interaction between the His-tagged SHANK3 SPN and KRASG12V recombinant proteins. MST measurements were performed using His-tag labelled SPN protein as a target and unlabelled KRASG12V as a ligand. The affinity curve and Kd-value are obtained from triplicate measurements. c, Isothermal titration calorimetry (ITC) titration and isotherms for the interaction between the active GMPPCP-form of KRASQ61H and wild-type (WT) SHANK3 SPN-ARR domain. Solid lines indicate fitting to the single-site-binding model at 25°C with 350 µM of KRASQ61H and 20 µM of SPN-ARR. d, ITC titration for the interactions between the inactive GDP-form of KRASQ61H and SHANK3 SPN-ARR. No interaction between the proteins is detected. e, Immunoprecipitation (IP) of wild-type SHANK3 (mRFP-SHANK3WT) and mutant KRAS (GFP-KRASG12V) after co-expression in HEK293 cells. The IPs and input lysates were analysed (blot) using anti-GFP and anti-SHANK3 antibodies, as indicated. Data are representative of three independent experiments. f, IP analyses of SHANK3 SPN wild-type and RAS-binding deficient mutant (GFP-SHANK3 SPN WT or R12E/K22D) co-expressed with mutant KRAS (KRASG12V-dsRed) in HEK293 cells and analysed using anti-GFP and anti-KRAS antibodies, as indicated. Data are representative of three independent experiments. g,h, FRET interaction between mCherry-tagged KRASG12V and GFP-tagged SHANK3 SPN WT (g) or RAS-binding deficient R12E/K22D mutant (h) in HEK293 cells. mCherry, control vector. Shown are the individual data points and the population average of each biological replicate (mean ± s.d.); n = 3 independent experiments. Unpaired Student’s t-test with Welch’s correction. i, Quantification of viable GFP positive MIA PaCa-2 cells expressing GFP-tagged WT or RAS-binding deficient (R12E/K22D) SHANK3 after silencing endogenous human *SHANK3* (36 h). Shown are individual data points and mean ± s.d. Data represents n = 4 independent experiments. One-way ANOVA with Holm-Sidak’s multiple comparison test.

To validate the SHANK3-KRAS protein-protein interaction in cells, we expressed fluorescently tagged KRASG12V and the full-length SHANK3 and performed pull-down experiments. Immunoblotting analyses showed that KRASG12V was co-immunoprecipitated from the cell lysates with the full-length SHANK3 (Fig. 2e). Based on its crystal structure (*29, 30*), the SHANK3 SPN domain contains the characteristic positively charged RAS-recognizing residues, R12 and K22, which are optimally positioned to form salt-bridges with the E37 and D38/Y40 residues of the KRAS Switch I region (residues from D30 to Y40; Fig. 2a). To analyse the interaction specificity between SHANK3 and KRAS, we introduced charge-reversing mutations into the R12 and K22 residues of SPN domain and co-expressed the SPN WT or SPN R12E/K22D domains with KRASG12V in cells. Pull-down analyses showed SHANK3 SPN domain co-precipitated KRASG12V from cells (Fig. 2f). In contrast, the R12E/K22D charge reversal double mutation in SHANK3 SPN abolished its association with KRASG12V (Fig. 2f). Accordingly, FRET-FLIM (Förster Resonance Energy Transfer by Fluorescence Lifetime Imaging Microscopy) of the GFP-tagged SHANK3 SPN WT, but not R12E/K22D mutant, demonstrated a significant protein-protein interaction with mCherry-KRASG12V in intact cells (Fig. 2g,f).

Taken together, these data demonstrate that SHANK3 directly interacts at low micromolar affinity with oncogenic KRAS, independent of the *KRAS* mutation itself, through conserved R12 and K22 residues characteristic of a RAS-effector interface.

To validate our hypothesis of SHANK3-KRAS interaction dependent tumour growth, we tested whether the WT SHANK3 re-expression, but not a KRAS-interaction deficient mutant, can rescue the growth inhibitory effect of *SHANK3* silencing. To test this, we chose *KRAS*-mutant MIA PaCa-2 pancreatic cancer cell line, as these cells were particularly sensitive to *SHANK3* siRNA-induced growth inhibition (Fig. 1a). We found that expression of siRNA resistant SHANK3 R12E/K22D mutant in *SHANK3*-silenced MIA PaCa-2 failed to rescue the cell death triggered by loss of SHANK3 (Fig. 2i and Extended Data Fig. 2e). In contrast, expression of siRNA resistant SHANK3 WT in the same cells restored cell viability back to control levels (Fig. 2i and Extended Data Fig. 2e). Thus, an intact KRAS-binding SPN domain is critical for the ability of SHANK3 to regulate *KRAS*-mutant cell survival.

### SHANK3 competes with RAF for KRAS binding and modulates downstream MAPK/ERK signalling

KRAS association with the plasma membrane and ability to recruit downstream effectors, such as RAF, are required for active KRAS signalling (*2, 35*). We observed that SHANK3 localization overlapped with mutant KRASG12V at the plasma membrane (Fig. 3a), prompting us to investigate the possibility of SHANK3 interacting with KRAS on the cell membrane.

**Figure 3.**
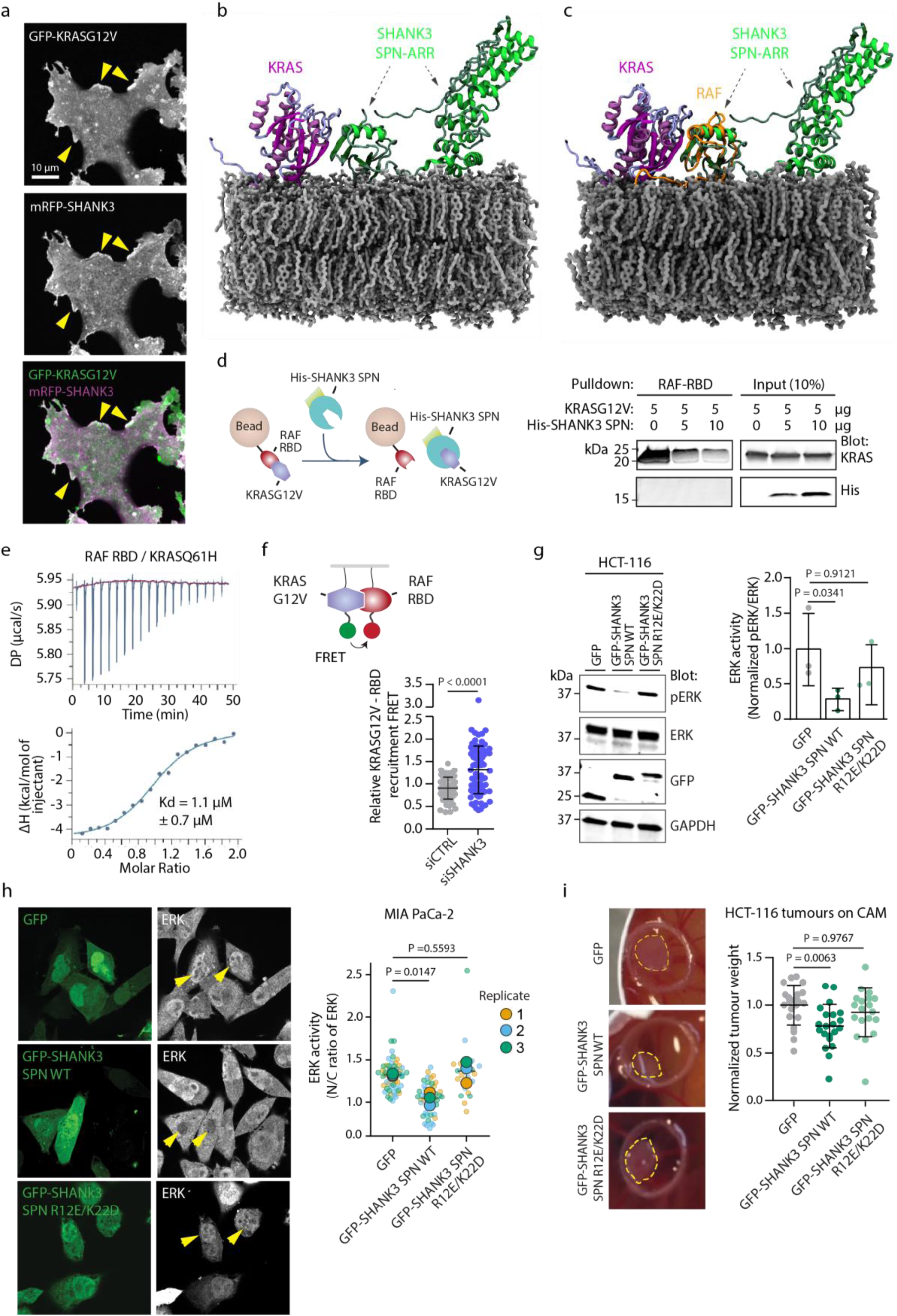
SHANK3 competes with RAF for the binding of active KRAS and limits downstream MAPK/ERK signalling. **a,** Microscopic co-localization of mutant KRAS (GFP-KRASG12V) and SHANK3 (mRFP-SHANK3) in cells. Yellow arrows point to co-localization sites at the membrane ruffles. Shown is a representative confocal slice (middle plane). b, KRAS— SHANK3 SPN-ARR in an open conformation modelled by aligning RBD and SPN domains of RAF and SHANK3 on the membrane composed of POPC (1-Palmitoyl-2-oleoyl-sn-glycero-3-phosphocholine)/ Phosphatidylinositol 4,5- bisphosphate/ Cholesterol). c, Structural alignment between KRAS—SHANK3 SPN-ARR (model) and nanodisc-bound KRAS—RAF complex (PDB:6PTW). d, Pulldown of RAF and SHANK3 competition for KRAS. Left: Principle of the bead competition assay. Right: RAF-RBD beads incubated with KRASG12V (5 µg) in presence or absence of His-SHANK3 SPN WT (0, 5 or 10 µg) were subjected to pulldowns. Input lysates and pulldown samples were analysed (blot) using anti- KRAS and anti-His antibodies, as indicated. Data are representative of three independent experiments. e, ITC titration and isotherms for the interaction between the active GMPPCP-form of KRASQ61H and RAF RBD. Solid lines indicate fitting to the single-site-binding model at 25°C with 20 µM of KRASQ61H and 200 µM of RAF-RBD in the same buffer conditions as in the Fig. 2c. Note that in the buffer used here the Kd value measured for KRASQ61H interaction with RAF RBD is 1.1 µM, which is somewhat higher than 0.1 µM reported for KRAS WT and 0.5 µM reported for KRASG13D. The main difference is the higher salt concentration in the buffer, which reduces the electrostatic contribution into the binding energy, with potentially a smaller effect from the mutation. f, KRASG12V-RBD-recruitment FRET assay in control or SHANK3 silenced cells. Right: A schematic representation of FRET assay. Left: Quantification of relative FRET efficiency between GFP-KRASG12V and mRFP-RAF-RBD co-expressed in control and SHANK3-silenced (smartpool SHANK3 siRNA) HEK293 cells. Shown are the individual data points and the mean ± s.d. from n = 78 - 86 cells from 3 independent experiments. Unpaired Student’s t-test with Welch’s correction. g,h, ERK1/2 signalling in *KRAS*-mutant cells expressing the KRAS interaction competent (GFP-SHANK3 SPN WT) or deficient (GFP-SHANK3 SPN R12E/K22D mutant) SHANK3 SPN domain. g, A representative immunoblot and quantification showing the ERK activation levels (phospho-ERK1/2 (Thr202/Y204) / total ERK relative to loading) in HCT116 (KRASG13D) cells. p-ERK, phospho-ERK1/2 (Thr202/Y204); ERK, total ERK; GAPDH, a loading control. Data represent the individual values and the mean ± s.d. (the mean of control is set to 1.0 by definition) from n = 3 independent experiments. Kruskal-Wallis test and Dunn’s post hoc test. h, Representative confocal images (the middle surface) and quantification of nuclear ERK (indicating ERK activity) in MIA PaCa-2 (KRASG12C) cells. Yellow arrowheads point to representative nuclei. N/C, nuclear to cytoplasmic ratio. Shown are the individual data points and the population average of each biological replicate (mean ± s.d.); n = 3 independent experiments. One-way ANOVA with Holm-Sidak’s multiple comparison test. i, In vivo CAM tumour growth in *KRAS*-mutant cells (HCT116) transiently expressing the KRAS interaction competent (GFP-SHANK3 SPN WT) or deficient (GFP-SHANK3 SPN R12E/K22D mutant) SHANK3 SPN domain. Shown are representative images and quantification. Tumours are within the yellow circles. Data are individual relative tumour weights and the mean ± s.d.; n = 19-21 tumours/group. Kruskal- Wallis test and Dunn’s post hoc test.

Based on the available SHANK3 structural data (SHANK3 SPN-ARR, (*29, 30*)), and our recent identification of the conformational opening of the SHANK3 SPN-ARR interface (*32*), we generated atomistic *in silico* models of the SHANK3 SPN and SPN-ARR fragments and simulated their binding to the plasma membrane (Extended Data Fig. 3a-d). These simulations indicated that SHANK3 SPN and ARR domains contain positively charged regions that could interact with the negatively charged plasma membrane. A recently published NMR structure defined a possible configuration of active KRAS binding to its downstream effector RAF on a lipid nanodisc membrane (*36*). Intrigued by the striking structural homology of the RAF RBD (Ras binding domain) and SHANK3 SPN domains (*29, 30, 36*) and our simulation data, indicative of a SHANK3 SPN-ARR membrane interaction, we generated a simulation model of the SHANK3 SPN-ARR-KRAS complex on the cell membrane (Fig. 3b). We observed that KRAS-bound SHANK3 (SPN-ARR domains) has evident interactions with the plasma membrane in its open configuration (Extended Data Fig. 3a-d; 32). An alignment between the SHANK3 SPN-ARR-KRAS model in an open conformation (Fig. 3b) and the nanodisc-bound KRAS-RAF (*36*) indicates strong overlap of SPN with the space occupied by the KRAS-interacting RAF (Fig. 3c and Extended Data Fig. 3a-d). These data imply that SHANK3 (SPN-ARR) competes with RAF for KRAS binding on the plasma membrane, possibly limiting KRAS downstream signalling.

To test this model, we first performed competition assays *in vitro*. Purified recombinant RAF-RBD and mutant KRASG12V proteins were incubated with increasing His-SHANK3 SPN protein concentrations followed by RAF-RBD pulldown (Fig. 3d). We observed a concentration-dependent reduction of KRAS binding to the RAF-RBD beads as a function of increasing His-SPN amounts (note that His-SPN and SPN-interacting KRAS are washed away in the pulldown) (Fig. 3d). ITC measurements for the interaction between active KRASQ61H and RAF-RBD (Fig. 3e; Kd = 1.1 μM ± 0.7 μM) and SHANK3 SPN-ARR (Fig. 2c; Kd = 5.4 μM ± 0.7 μM) under the same buffer conditions showed relatively small difference in affinities, indicating a possibility for SHANK3 to compete with RAF-RBD for KRAS binding.

We then examined whether KRAS effector recruitment was affected in the absence of SHANK3. Silencing of endogenous *SHANK3* enhanced RAF-RBD and KRASG12V interaction in cells, as determined using an established cell-based FRET assay (*37*) for KRAS effector recruitment and downstream signalling (Fig. 3f). Collectively, these data indicate that SHANK3 can effectively compete with RAF to interact with mutant KRAS and thus may be able to influence KRAS downstream signalling in cells.

The MAPK/ERK pathway (also known as the RAS-RAF-MEK-ERK pathway) is a critical signalling node in *KRAS*-mutant cancers. Active RAS mutants recruit RAF to signal through this pathway to induce ERK phosphorylation and nuclear translocation and to promote ERK-dependent cell proliferation (*8, 9*). We thus sought to test whether the ability of SHANK3 to compete with RAF for active KRAS binding could subsequently modulate downstream MAPK/ERK signalling. Transient overexpression of SHANK3 SPN in *KRAS*-mutant HCT116 cells significantly decreased ERK1/2 phosphorylation (Fig. 3g) and diminished ERK1/2 nuclear translocation in *KRAS*-mutant MIA Paca-2 cells (Fig. 3h). The RAS-binding deficient SPN mutant (R12E/K22D), in contrast, did not suppress ERK1/2 phosphorylation (Fig. 3g) and had no significant effect on ERK1/2 translocation to the nucleus (Fig. 3h). In line with SHANK3 SPN-mediated attenuation of ERK-activity, SHANK3 SPN WT overexpression, but not SPN R12E/K22D, in HCT116 cells restrained KRAS-driven tumourigenic growth in the chick embryo CAM CRC xenografts (Fig. 3i). These data demonstrate that SHANK3 SPN competes with RAF for active KRAS binding and limits oncogenic signalling via the MAPK/ERK pathway.

### Loss of SHANK3 triggers cell death through RAS-MAPK pathway hyperactivation

Recent studies indicate that the level of MAPK/ERK activity in tumour cells needs to be carefully maintained within a precise range: the signalling has to be sufficiently high to support tumour growth and yet below the toxic level that triggers apoptosis or senescence (*38–40*).

We observed that *SHANK3*-silenced *KRAS*-mutant pancreatic (PANC-1) and lung (A549) cancer cells showed a marked 8‒30-fold increase in ERK1/2 phosphorylation (Fig. 4a). In contrast, AKT activity was highly variable but not significantly changed (Extended Data Fig. 4). To further validate these changes in ERK activity in living *SHANK3*-silenced cells on the single cell level we used an ERK-KTR (Kinase Translocation Reporter) (*41*), which shuttles from the cytoplasm to the nucleus in response to changes in ERK activation state (Fig. 4b). *SHANK3* silencing in PANC-1 cells significantly increased the cytoplasmic-to-nuclear (C/N) ratio of the KTR, further demonstrating that SHANK3 depletion induces MAPK/ERK signalling overactivation in *KRAS*-mutant cells (Fig. 4b).

**Figure 4.**
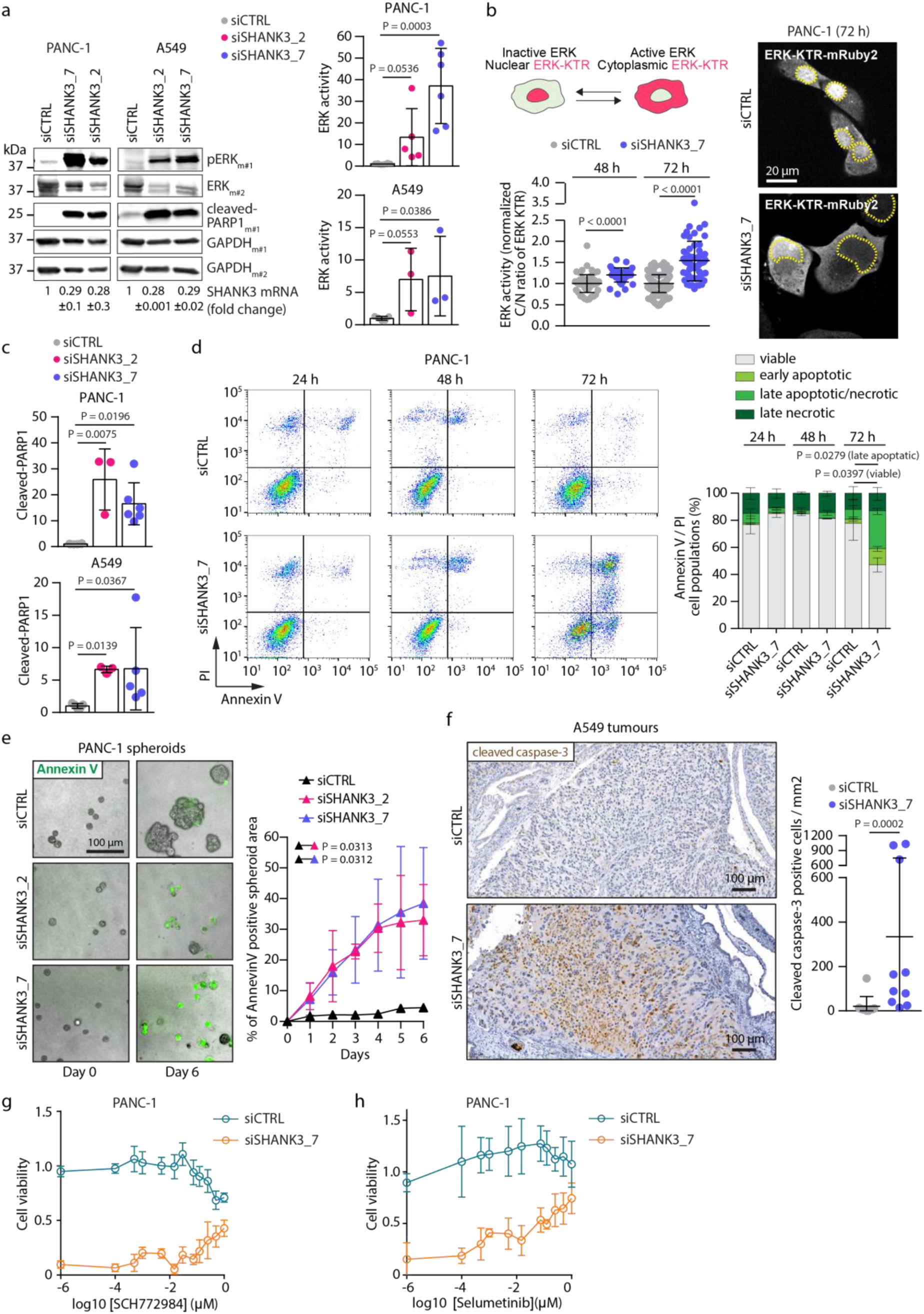
SHANK3 depletion triggers RAS-MAPK pathway hyperactivation and apoptosis in *KRAS*-mutant cells. **a,** ERK signalling in *SHANK3*-silenced *KRAS*-mutant in PANC-1 (KRASG12D) and A549 (KRASG12S) cells. Left: Representative immunoblots of the indicated proteins in control or *SHANK3*-silenced cells analysed three days after silencing. p-ERK, phospho-ERK1/2 (Thr202/Y204); ERK, total ERK; cleaved-PARP1, indicative of apoptosis; GAPDH, a loading control; m#1 and m#2, duplicate membranes. Below the immunoblots: *SHANK3* mRNA levels (fold change) showing the efficiency of *SHANK3* silencing. Right: Quantification of ERK1/2 activity (phospho-ERK1/2 (Thr202/Y204) / total ERK relative to loading) in control or *SHANK3*-silenced cells. Data points are the individual experiments and bars are the mean±s.d.; n = 3-6 independent experiments. Kruskal-Wallis test and Dunn’s post hoc test. b, ERK activation by *SHANK3* silencing. Top: A schematic representation of the ERK-KTR biosensor (red) that translocates from the nucleus to the cytoplasm upon ERK activation. Right: Representative confocal images of control and SHANK3-silenced PANC1 cells stably expressing ERK- KTR-mRuby2 three days after silencing. Nuclei outlined by yellow dashed line. Left: Quantification of cytoplasmic/nuclear (C/N) ratio of ERK-KTR analysed two and three days after silencing. Data show individual cells and the population average of each biological replicate (mean±s.d.); n = 3 independent experiments. One-way ANOVA with Holm-Sidak’s multiple comparison test. c, Immunoblotting analyses of PARP1 cleavage (apoptosis) in control or *SHANK3*-silenced *KRAS*-mutant PANC-1 and A549 cells analysed three days after silencing. Shown are the individual values and mean±s.d. (the mean of control is set to 1.0 by definition; n = 3-6 independent experiments). Kruskal-Wallis test and Dunn’s post hoc test. Representative immunoblots are shown in 4a. d,e, Cell death in control or *SHANK3*-silenced PANC-1 cells/spheroids. d, Representative scatter plots and quantification of AnnexinV-FITC / PI flow cytometry data from 2D-cultured cells 1, 2 and 3 days after silencing (mean ± s.d.; n = 5 independent experiments). Unpaired Student’s t-test with Welch’s correction. e, Representative images and analysis of apoptosis (AnnexinV positive area) in spheroids grown in 3D Matrigel (mean ± s.d.; n = 3 independent experiments). One-way ANOVA with Holm-Sidak’s multiple comparison test (at the endpoint). f, Cleaved caspase-3 levels in the CAM tumours. Control and *SHANK3*-silenced tumours were inoculated on CAM as in Fig. 1e-f. Shown are representative images and quantitative analysis of A549 tumours (mean ± s.d.; n = 10 tumours; from *SHANK3*-silenced PANC-1 not enough residual tumour was available for meaningful analyses). Mann Whitney test. g,h, ERK/MEK-dependence of SHANK3-induced cell death in *KRAS*-mutant cells. Cell viability of control or *SHANK3*-silenced PANC-1 cells treated with the indicated concentrations of ERK inhibitor SCH772984 (g) or MEK inhibitor selumetinib (h). Shown is mean±s.d; n = 3-4 measurements (representative graph from 2 independent experiments).

In addition to ERK hyperactivation, *SHANK3*-silencing significantly increased the levels of cleaved- PARP1 in PANC-1 and A549 cells (Fig. 4c), and the number of AnnexinV/PI-positive PANC1 cells (Fig. 4d). *SHANK3* silencing in PANC1 cells impaired formation/growth of proper 3D spheroids in Matrigel (Fig. 1d) and significantly increased numbers of AnnexinV-positive apoptotic cells over time (Fig. 4e). We also detected notably higher cleaved caspase-3 staining in *SHANK3*-silenced *KRAS*-mutant A549 CAM tumours compared to control tumours (Fig. 4f). These data indicate that loss of SHANK3 induces apoptosis in *KRAS*-mutant cells.

To determine whether the cytotoxic effects of *SHANK3*-silencing in *KRAS*-mutant cells depend on the MAPK/ERK pathway, we treated PANC-1 cells with low doses of the ERK inhibitor SCH772984 and MEK inhibitor selumetinib. In control silenced cells, low doses of the ERK and MEK inhibition had no/modest effect on cell viability cells (Fig. 4g,h), as previous described (*42*). In *SHANK3*-silenced cells, both SCH772984 and selumetinib counter-acted the proliferation defect in a dose-dependent manner (Fig. 4g,h). These results indicate that the anti-proliferative effects and reduced cell viability triggered by the depletion of SHANK3 in *KRAS*-mutant cells are linked to enhanced MAPK/ERK activity.

### SHANK3 depletion impairs the growth of established *KRAS*-mutant tumours

To evaluate the requirement of SHANK3 in maintaining the growth of established tumours, we generated PANC-1 cell clones with a doxycycline (Dox)-inducible shRNA against *SHANK3* (Fig. 5a). We first verified that dox induced loss of SHANK3 protein and suppressed of the growth of these lines *in vitro* (Fig. 5a-e and Extended Data Figs. 5 and 6). Consistent with the results obtained by siRNA-mediated silencing, inducible silencing of *SHANK3* (using a shRNA targeting sequence distinct from the two siRNAs) strongly increased ERK phosphorylation and consequent PARP1 cleavage in a time-dependent fashion (Fig. 5b and Extended Data Fig. 5). The induction of *SHANK3*-silencing also dampened the growth of established 3D spheroids and was accompanied by a significant increase in AnnexinV-positive regions within the spheroids over time (Fig. 5c-e and Extended Data Fig. 6).

**Figure 5.**
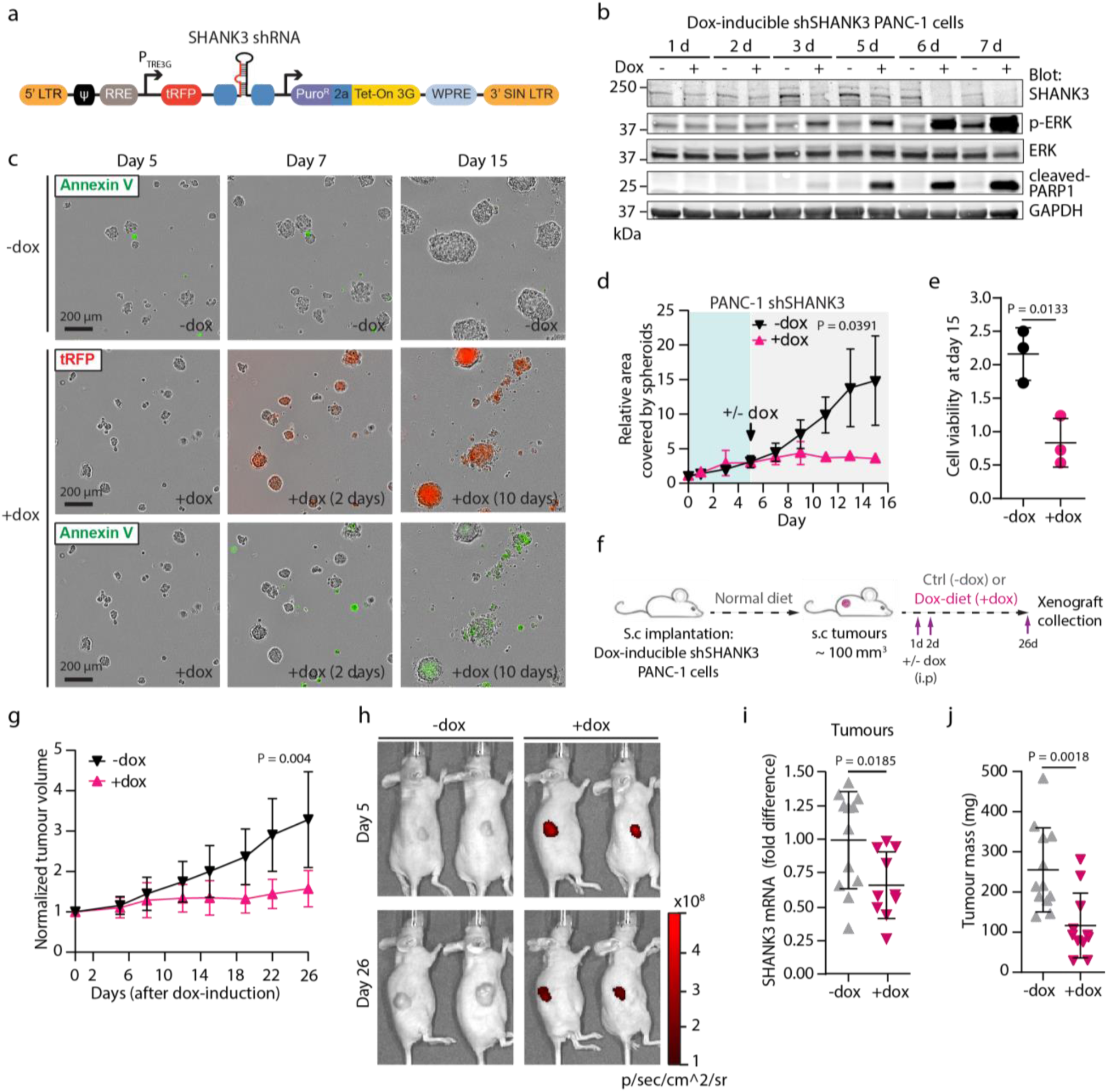
SHANK3 depletion impairs the growth of pre-existing *KRAS*-mutant PDAC tumours. **a,** A schematic representation of the lentiviral vector for tetracycline/doxycycline (Dox)-inducible synthesis of *SHANK3* shRNA. PTRE3G inducible promoter with tetracycline response element is activated by the Tet-On® 3G transactivator protein in the presence of doxycycline. 5’LTR, 5’long terminal repeat; Ψ, Psi packaging sequence; tRFP, TurboRFP reporter for visual tracking expression upon doxycycline induction; PuroR, puromycin resistance gene; 2a, self-cleaving peptide enables the expression of both PuroR and Tet-On® 3G transactivator from a single RNA pol II promoter; Tet-On® 3G encodes the doxycycline-regulated transactivator protein, which binds to PTRE3G promoter only in the presence of doxycycline; WPRE, Woodchuck Hepatitis Post-transcriptional Regulatory Element; 3’ SIN LTR, 3’ Self-inactivating Long Terminal Repeat. b, ERK activation kinetics over the course of *SHANK3* silencing in *KRAS*-mutant cells. Representative immunoblots showing SHANK3, p-ERK (indicative of ERK activation) and cleaved-PARP1 (indicative of apoptosis) levels in control (- dox; non-induced) and doxycycline-induced (+ dox) sh*SHANK3*-expressing PANC-1 cells (mix of two independent clones) collected at different time points (days 1-7). GAPDH serves as a loading control. c-d, Growth and viability of control (- dox) and doxycycline-induced (+dox) sh*SHANK3*-expressing PANC-1 spheroids. SHANK3-depletion was induced in established spheroids by doxycycline (+dox) at day 5 and continued until day 15 (i.e. 10 days after dox-induction). c, Representative images. SHANK3 depletion was observed by tRFP reporter (visual tracking of dox-induction and *SHANK3* shRNA expression). Apoptotic AnnexinV positive cells are shown in green. d, Quantification of spheroid growth. e, Colorimetric assay for cell viability in spheroids measured at day 15 (the endpoint). Data are the mean±s.d. from n= 3 independent experiments (d,e). Unpaired Student’s t-test with Welch’s correction (at the endpoint). f-j, Growth of established tumours in mice after SHANK3 depletion. f, The experimental outline with the PANC-1 pancreatic tumour cells stably expressing dox-inducible *SHANK3* shRNA vector and tRFP reporter to validate shRNA induction. When palpable tumours appeared, mice received either dox-containing (+ dox) or normal diet (- dox) (i.p admistration of dox or PBS at day 1 and 2). g, Tumour volumes after starting the dox-treatment (normalised to tumour volumes at the start of the dox induction). h, Representative IVIS images of tRFP reporter expression in tumours 5 and 26 days after dox- induction. i, *SHANK3* gene expression (mRNA levels) in tumours with doxycycline-inducible SHANK3 knockdown (+dox) at the end of the experiment. j, Tumour weights at the end of the experiment (26 days after dox-induction). Data represent individual tumours and the mean ± s.d.; n = 11 (+ dox) and 12 (- dox) tumours (g,i,j). Unpaired Student’s t-test with Welch’s correction.

**Figure 6.**
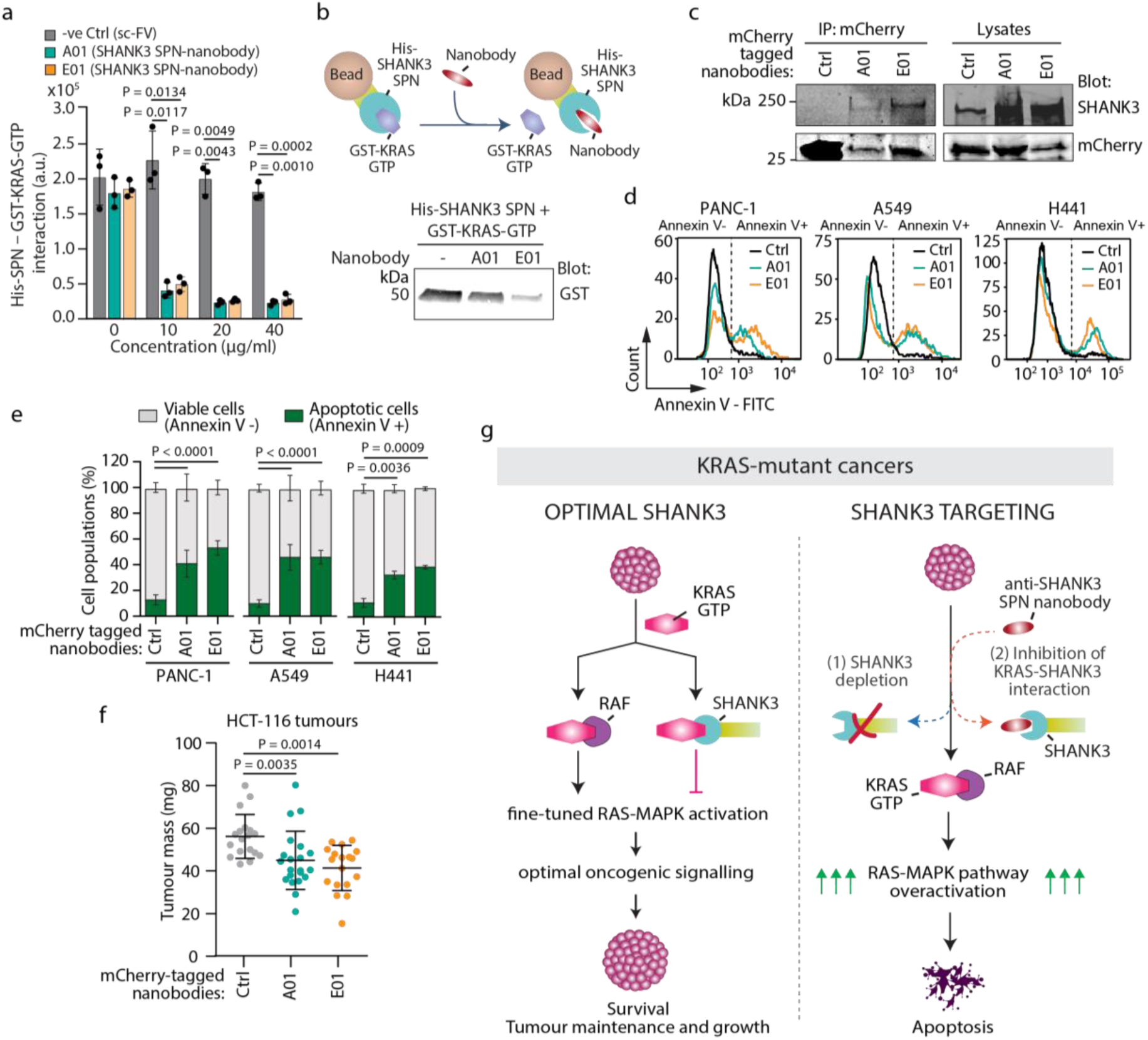
Targeting the KRAS-SHANK3 interaction by anti-SHANK3 nanobodies induces apoptosis and inhibits KRAS-driven tumour growth *in vivo.* **a-c,** Inhibitory function of anti-SHANK3 nanobodies on KRAS – SHANK3 interaction. a, The effect of two independent SHANK3 SPN nanobodies (A01 and E01) at the indicated concentration on the binding of His-tagged SHANK3 SPN to GST-tagged KRAS-GTP as measured by ELISA. Data are mean ± s.d. of the Eu-signal; n = 3 independent experiments. b, Top: The experimental outline of His-SHANK3 SPN pulldown of GST-KRAS-GTP in the presence or absence of anti-SHANK3 SPN nanobodies. Bottom: A representative blot from 2 independent experiments. c, Specificity analysis of anti-SHANK3 SPN nanobodies. mCherry-tagged anti-SHANK3 SPN (A01, E01) and control nanobodies expressed in A549 cells were immunoprecipitated (IP) from cell lysates and blotted as indicated. A representative blot from 2 independent experiments. d,e, Cell viability by anti-SHANK3 nanobodies. Representative histograms (d) and quantification (e) of apoptosis in control or anti-SHANK3 SPN-nanobodies expressing PANC-1, A549 and H441 cells 4 days after transfections using AnnexinV-FITC flow cytometry assay. Data represent mean ± s.d.; n = 3-4 independent experiments/cell type. Two-way ANOVA and Sidak’s post hoc test. f, In vivo tumour growth on CAM membranes with HCT116 cells transiently expressing mCherry-tagged anti-SHANK3 SPN (A01 and E01) or control nanobodies. Data represent individual tumours and the mean ± s.d.; n = 18-21 tumours/treatment group. Kruskal-Wallis test and Dunn’s post hoc test. g, Schematic model of SHANK3-controlled cell fate in *KRAS*-mutant cancers. SHANK3 directly interacts with KRAS and competes with RAF for KRAS binding to sustain oncogenic RAS-MAPK/ERK signalling at an optimal level (i.e. below toxic oncogenic signalling) in *KRAS*-mutant cancers. SHANK3 silencing (i) and inhibition of SHANK3-KRAS interaction (ii) drive *KRAS*-mutant cells into cell death.

To study the effect of Dox-induced SHANK3-depletion in established tumours *in vivo*, we implanted sh*SHANK3* expressing cells into mice, allowed them to form tumours (∼100 mm^3^) and thereafter subjected the mice to Dox-supplemented or normal diet (Fig. 5f). shRNA induction was monitored by imaging red fluorescence (tRFP reporter) and tumour growth was followed by bi-weekly palpations. SHANK3 depletion in the established tumours led to robust inhibition of tumour growth when compared to control tumours (Fig. 5g,h and Extended Data Fig. 7). At the end of the experiment, *SHANK3* mRNA expression was still significantly suppressed in Dox-induced tumours (Fig. 5i). The reduced growth of established *SHANK3*-depleted tumours was recapitulated when measuring the tumour masses at the end of the experiment (Fig 5j). These *in vivo* data highlight the potential of SHANK3 targeting as an anti-cancer therapy in pre-existing *KRAS*-mutant tumours.

### Nanobodies disrupting the SHANK3-KRAS interaction drive *KRAS*-mutant cells into apoptosis

As an initial step towards the development of a SHANK3-based therapy, we generated nanobodies that interfere with the KRAS-SHANK3 interaction and assessed their efficacy in *KRAS*-mutant cancer cells. A phage display library screen identified two distinct single-domain antibody fragments (VHH-binders; nanobodies A01 and E01) directed against the SHANK3 SPN domain. Both nanobodies robustly inhibited SHANK3-KRAS-GTP interaction in ELISA-based binding assays as well as in *in vitro* pull-down assays (Fig 6a and b). When expressed in cells, both anti-SHANK3 SPN nanobodies co-precipitated endogenous SHANK3 (Fig. 6c). In functional viability studies with *KRAS*-mutant pancreatic (PANC-1) and lung (A549 and H441) cancer cells, both nanobodies increased the number of Annexin V^+^ apoptotic cells in all three cell lines (Fig. 6d and e). Finally, while control nanobody-expressing cells rapidly formed tumours in the CAM xenograft model, overexpression of A01 and E01 anti-SHANK3 nanobodies significantly reduced the KRAS-driven tumour growth *in vivo* (Fig. 6f). These data indicate that disrupting SHANK3 interaction with KRAS results in a similar loss of cell viability and apoptosis induction in *KRAS*-mutant cells as with RNAi-mediated SHANK3 depletion (Fig. 6 g).

Together, these findings support the concept that hyperactivating RAS-MAPK pathway in *KRAS*-mutant cells, by ablating the SHANK3-KRAS interaction, could have a therapeutic impact on *KRAS*-mutant cancers.

## Discussion

Our work identified the SHANK3 scaffold protein as an essential regulator of active and mutant KRAS. SHANK3 directly interacts with KRAS, competes with RAF for KRAS binding on the plasma membrane and sets downstream MAPK/ERK signalling to an optimal level to sustain proliferative capacity and limit active ERK levels from reaching a lethal signalling threshold (Fig. 6g). We demonstrate that by disrupting the SHANK3-KRAS interaction, thus removing an endogenous KRAS signalling brake, we can trigger a cytotoxic level of ERK activity that results in reduced cell proliferation, apoptosis induction and impaired tumour growth in *KRAS*-mutant xenograft models. This indicates that KRAS-driven cancer cells require intermediate levels of SHANK3-KRAS association to support tumourigenic growth.

Targeting SHANK3 to induce RAS-MAPK hyperactivation-induced apoptosis represents a conceptually novel therapeutic approach for treating *KRAS*-mutant cancers. Firstly, we show that SHANK3 binding to oncogenic KRAS is not dependent on specific *KRAS*-activating mutations. Therefore, targeting the SHANK3- KRAS interaction represents a new pan-KRAS-mutant compatible strategy for selective killing of *KRAS*-mutant cancer cells. Secondly, current efforts strive to develop KRAS inhibitors (*12, 15–17, 43*), whereas we describe a mechanism to hyperactivate KRAS-MAPK signalling to cytotoxic levels by disrupting KRAS interaction with SHANK3.

We provide proof-of-concept evidence of the ability of inhibitors targeting the KRAS-SHANK3 interaction to trigger apoptosis and limit the growth of *KRAS*-mutant cancers. We foresee SHANK3 as a strong candidate for innovative drug discovery programs to target a broad spectrum of *KRAS*-mutant cancers (*e.g.* 95% of PDAC; ∼30% of NSCLC) (*3–5, 20*). Although targeting intracellular proteins by antibodies has been difficult, the use of nanobodies with nanoparticles and cell-penetrating peptides offers new opportunities (*44, 45*). Furthermore, development of small molecule inhibitors of protein-protein interactions has recently become possible in pharma industry, as exemplified by the successful generation of BH3-mimetics (*46*). Similarly, several siRNA therapeutics have already gained FDA approval (*47–49*). Our findings that *KRAS*-mutant tumour growth can be inhibited in *in vivo* settings either by modulating *SHANK3* RNA expression or by blocking KRAS-SHANK3 protein-protein interaction by nanobodies with a reasonable efficacy should offer multiple options for further pharmaceutical development.

In summary, we have discovered here that disruption of the SHANK3-KRAS interaction in different *KRAS*-mutant cancers effectively hijacks the tumour’s own oncogenic signalling machinery and drives it to kill itself. Therefore, SHANK3 emerges as an attractive new therapeutic target in the field aiming to address the huge unmet need for effective pan-KRAS therapy.

## Methods

### Cell lines and cell culture

PANC-1, AsPC-1, SW1990, PaTu8902, MIA PaCa-2 (human pancreatic ductal adenocarcinoma cell lines, all from ATCC), A549 (human lung adenocarcinoma, ATCC), HCT-116 (human colorectal carcinoma, ATCC), HT-29 (human colorectal adenocarcinoma, ATCC), HEK293 (human embryonic kidney, ATCC) and U2OS (osteosarcoma, ATCC) cells were grown in DMEM (Dulbecco’s modified Eagle’s medium, Sigma-Aldrich) supplemented with 10% FBS and 2 mM L-glutamine.

Su86.86 (human pancreatic adenocarcinoma, ATCC), YAPC (human pancreatic carcinoma, DSMZ), H441 (human lung adenocarcinoma, ATCC), HCT-15 (human colorectal adenocarcinoma, ATCC), H292 (human lung carcinoma, ATCC), H226 (human lung squamous cell carcinoma, ATCC) and H266 (human non-small cell lung carcinoma) cells were cultured in RPMI-1640 medium supplemented with 10 % FBS and 2mM L-glutamine.

Panc10.05 (human pancreatic adenocarcinoma, ATCC) cells were cultured in RPMI-1640 medium supplemented with 10 % FBS, 2mM L-glutamine and 10 Units/ml human recombinant insulin. ARPE-19 (human retinal pigmented epithelium, ATCC) cells were grown in DMEM:F12 (Gibco) supplemented with 10% FBS and 2 mM L-glutamine.

Cells were regularly tested by MycoAlert^TM^ Mycoplasma Detection Kit (Cat. no. LT07-418, Lonza) with MycoAlert^TM^ Assay Control Set (Cat. no. LT07-518, Lonza) and found to be free from mycoplasma contamination. Cell lines used in this study were authenticated by DSMZ.

### siRNAs and DNA constructs

The siRNAs targeting human *SHANK3* were SMARTpool ON-TARGETplus Human SHANK3 siRNA (Cat. no. L- 024645-00, Dharmacon), individual Human SHANK3 siRNA_2 (Cat. no. S100717710 Hs_SHANK3_2 siRNA, Qiagen) and individual ON-TARGETplus Human SHANK3 siRNA_7 (Cat. no. J-024645-07 Dharmacon). The control siRNA was Allstars negative control siRNA (Cat. no. 1027281, Qiagen).

EGFP-tagged SPN domain and mRFP-tagged SHANK3 has been described in (*29*). pHAGE-EGFP-Shank3 (GFP-SHANK3 WT) was kindly supplied by Alex Shcheglovitov. The R12E/K22D mutation was introduced by site directed mutagenesis (Gene Universal). pmGFP-KRASG12V and mRFP-RBD with the RAS binding domain (RBD) of CRAF have been described earlier (*50, 51*). pmCherry-KRASG12V construct was generated by replacing pmGFP from pmGFP-KRASG12V with pmCherry from pmCherry-C1 vector (Clontech Laboratories Inc.) using NheI and BsrGI restriction sites. DsRed-KRASG12V was a gift from J Lorens. His6-KRASQ61H (Plasmid #25153, Addgene) was a gift from Cheryl Arrowsmith. pLentiPGK-Blast-DEST-ERKKTRmRuby2 (Plasmid #90231, Addgene) was a kind gift from Markus Cover (*41*). mCherry-tagged anti-SHANK3-SPN nanobodies A01 and E01 were generated by Hybrigenics. In addition, peGFP-C1 and pHAGE-CMV-eGFP-W (PlasmID, Harvard Medical School) and pmCherry-C1 were used as controls.

### Transient transfections

Lipofectamine 3000 and P3000™ Enhancer Reagent (Cat. no. L3000001, Thermo Fisher Scientific) or jetPRIME (Cat. no. 101000046, Polyplus) were used to transiently transfect cells with plasmids according to the manufacturer’s protocol and the cells were cultured for 24 h. SiRNA silencing was performed using 30-67 nM siRNA (siRNA targeting *SHANK3* or negative control siRNA) and Lipofectamine^®^ RNAiMAX Reagent (Cat. no. 13778075, Thermo Fisher Scientific) according to the manufacturer’s protocol, and cells were cultured for 24 h. One day post-transfections medium was changed to full culture medium and cells were used for the experiments at the indicated time points.

### Proliferation assay

Cells were seeded on a 96-well plate and transfected with siRNAs on the following day, as described above. To perform a proliferation screen in multiple cancer cell lines 5000-10000 cells were seeded in the 96-wells depending on the growth rate of control cells. Proliferation was measured for 4 days using the IncuCyte S3 Live-Cell Analysis system (10x objective). Wells were imaged every two hours (brightfield and green phase; acquisition time 300 ms). Culture medium was changed every 2-3 days. Analysis was performed using IncuCyte S3 software. The analysis definition was set using the following parameters: segmentation (background-cells), clean-up (hole fill), filters (area, eccentricity, mean intensity, integrated intensity). A mask was set to the best fit of cell confluence to quantify the area covered by cells. Normalised proliferation was calculated from time-lapse imaging by dividing the area covered by cells at every time-point by the area of the first time point (t = 0; averaged reading used from replicate wells).

### Colony formation

Cells were siRNA silenced for 24 h as described above. A day post-silencing, 250 cells were seeded per well on a 6-well plate in full medium. The culture medium was changed every 2-3 days and the assay was ended on day 10-14 (PANC-1, A549 cells). Cell colonies were fixed with 4 % paraformaldehyde (PFA) in phosphate-buffered saline (PBS) for 15 min and washed with PBS. Then, colonies were stained with 0.2 % crystal violet in 10 % EtOH for 10 min at room temperature (RT) and washed with PBS. Plates were scanned and analysed using a Colony area ImageJ plugin previously described by Guzmán et al. 2014 (*52*).

### 3D spheroid formation assay

3D spheroid formation was measured by a previously described method (*53*) where cells are embedded between two layers of Matrigel. The inner wells of an angiogenesis 96-well µ-plate (Cat. no. 89646, Ibidi GmbH) were coated with 10 µl of 50% Matrigel (diluted in full cell culture medium; Matrigel stock 9 mg/ml, Cat. no. 354230, Corning), centrifuged for 20 min at 200 g (4°C) and incubated for 1 hour at 37°C. Next, the upper wells were filled with 20 µl of cell suspension in 25% Matrigel (500 cells/ well), centrifuged for 10 min at 100 g and incubated at 37°C for 4h or overnight. Then, wells were filled with full cell culture medium and spheroid formation was measured for 6-15 days with IncuCyte S3 Live-Cell Analysis system (10x objective). Wells were imaged every two hours (Phase + brightfield and green phase, acquisition time 300 ms). Culture medium was changed every 2-3 days. Analysis was performed using IncuCyte S3 software. Analysis definition was set using the following parameters: segmentation (background-cells), clean-up (hole fill); filters (area, eccentricity, mean intensity, integrated intensity). A mask was set to the best fit of cell confluence to quantify cell/spheroid area.

### Quantitative real-time polymerase chain reaction

Total cellular RNA was extracted using the NucleoSpin^®^ RNA kit (Cat. no. 740955.50 Macherey- Nagel) and 1 µg of the extracted RNA was used as a template for cDNA synthesis by high-capacity cDNA reverse transcription kit (Cat. no. 4368814, Applied Biosystems) according to the manufacturer’s protocol. Tumours were homogenised with T 25 ULTRA-TURRAX® (Ika) and total RNA was extracted using TRIsure™ (Cat. no. BIO-38032 Bioline Ltd). Expression levels of *SHANK3* were determined by TaqMan® qRT-PCR reaction using QuantStudio™ 12K Flex Real-Time PCR System (Thermo Fisher Scientific). The level of glyceraldehyde 3-phosphate dehydrogenase (*GAPDH*) expression was used as a reference (endogenous control). Taqman® Universal Master Mix II included necessary components for qRT-PCR reaction (Cat. no. 4440040, Thermo Fisher Scientific). TaqMan® Gene Expression Assays (Thermo Fisher Scientific) were used to detect *SHANK3* (Assay ID: Hs00873185_m1) and *GAPDH* (Assay ID: Hs02786624_g1). Relative expression was calculated by a comparative CT (ΔΔCT) method using the expression level of *GAPDH* as a reference for the quantification.

### Immunoblotting

Cells were transfected with siRNA or plasmid as described above. Cells were collected into lysis buffer [50 mM Tris-HCl (pH 7.5), 150 mM NaCl, 0.5% Triton-X, 0.5% glycerol, 1% SDS, complete protease inhibitor (Sigma-Aldrich) and PhosSTOP (Sigma-Aldrich)] and protein extracts were sonicated. Protein levels were measured by Bio-Rad protein quantification kit. Sample buffer was added and samples were boiled for 5 min at 95°C. Proteins were then separated using SDS-PAGE under denaturing conditions (4–20% Mini-PROTEAN TGX Gels) and transferred onto nitrocellulose membranes by semi-dry turbo blot (BioRad Laboratories). Membranes were blocked with 5 % bovine serum albumin (BSA) in TBST (Tris-buffered saline and 0.1% Tween 20) for 1 hour at RT. Primary antibodies were diluted in 5 % BSA in TBST and incubated with membranes overnight at +4°C. After primary antibody incubation, membranes were washed three times with TBST for 5 min at RT. Fluorophore-conjugated or ECL HRP-linked secondary antibodies (GE Healthcare) were diluted 1:5000 in 5 % BSA in TBST or in blocking buffer (Cat. no. 37538, Thermo Fisher Scientific) in PBS (1:1) and were incubated with membranes for an hour at RT. Membranes were scanned using an infrared imaging system (Odyssey; LI-COR Biosciences) or using ECL Plus Western blotting reagent (Cat. no. RPN2232, GE Healthcare) and film. Band intensity was determined using Fiji (ImageJ; National Institutes of Health) or Image Studio Lite (LI-COR).

The following primary antibodies were used: SHANK3 (Cat. No. HPA003446, Atlas antibodies and Cat. no. sc-30193, Santa Cruz), GFP (Cat. no. ab1218, Abcam), KRAS (Cat. no. WH0003845M1, Sigma-Aldrich), GAPDH (Cat. no. 5G4-6C5, Hytest), HSP70 (Hsc70/Hsp73; Cat. no. ADI-SPA-815, Enzo), phopho-ERK1/2 (Thr202/Tyr204) (Cat. no. 4370S, Cell Signaling), ERK1/2 (Cat. no. 91025, Cell Signaling), phospho-AKT (Ser473) (Cat. no. 9271, Cell Signaling), AKT (Cat. no. 9272, Cell Signaling) and cleaved-PARP1 (Cat. no. ab4830, Abcam). All primary antibodies were used at 1:1000 dilution, except the SHANK3 antibody which was used at 1:100 (Cat. no. sc-30193, Santa Cruz) or 1:500 (Cat. No. HPA003446, Atlas antibodies) dilution.

### Co-immunoprecipitations

HEK293 cells were transiently transfected with (1) mRFP-tagged SHANK3 WT together with GFP-tagged KRASG12V or control (GFP only); or (2) GFP-tagged SHANK3 SPN WT, SHANK3 SPN R12E/K22D or control (GFP only) together with dsRed-tagged KRASG12V. A549 cells were transiently transfected with control plasmid (pmCherry) or the nanobodies (pmcherry-A01 or pmcherry-E01). 24 hours post-transfection the cells were lysed using IP-lysis buffer (40 mM Hepes-NaOH, 75 mM NaCl, 2 mM MgCl2, 1% NP40, protease and phosphatase inhibitors), cleared by centrifugation, and subjected to immunoprecipitation of RFP/mCherry/dsRed-tagged or GFP-tagged fusion proteins using RFP-trap or GFP-trap matrix (Chromotek), respectively. Input and precipitate samples were analysed by immunoblotting.

### Production of recombinant KRAS mutants, SHANK3 fragments and BRAF

All proteins were produced in transfected *E. coli* BL21 cultures using IPTG induction. Recombinant KRASG12V protein was generated as a glutathione S-transferase (GST) fusion protein, purified using gluthathione agarose and finally the GST was cleaved off. To produce recombinant KRASQ61H, we expressed the plasmid containing human His6-KRASQ61H activating oncogenic mutant (residues 1-169) in *E. coli*, purified the protein using Ni-resin and exchanged the bound nucleotide for non-hydrolysable GTP analogue GMPPCP using alkaline phosphatase beads (Sigma-Aldrich) and following the protocol of John et.al. (*54*). Bacterially produced His-tagged RBD domain of human BRAF was purified using Ni-columns and finally the His-tag was cleaved off using protease. His-tagged SPN fragment of SHANK3 was also produced, and purified using Ni-resins. His-tagged SPN-ARR fragment was produced and purified as previously described (*32*). The homodispersity of all proteins was verified with sodium dodecyl sulphate polyacrylamide gel electrophoresis (SDS-PAGE). The detailed methodology for production and purification of the recombinant proteins is described in the Extended Data Methods.

### Microscale thermophoresis (MST)

Interaction of recombinant SPN fragment of SHANK3 and recombinant KRASG12V was measured using MST. His-SPN was labelled using Monolith His-Tag Labelling Kit Red-tris-NTA, fluorescent dye (Cat no. L008, NanoTemper Technologies) and applied at the final concentration of 50 nM in His-SPN SEC buffer having 0.05% Tween-20. A 12-point two-fold dilution series of unlabelled KRASG12V was mixed with labelled His-SPN protein. The final concentration of KRASG12V protein range from 200 µM to 200 nM (KRASG12V). Microscale thermophoresis (MST) experiments were conducted in triplicate using Monolith automated capillaries (Cat no. MO-AK002, NanoTemper Technologies) with a Monolith NT Automated system (NanoTemper Technologies GmbH, Munich) to determine the binding affinity between His-SPN and KRASG12V. The dissociation constant was then calculated using a single-site binding model to fit the curve and errors with standard error of mean (SEM) using GraphPad Prism version 8.4.2 for Windows (GraphPad Software Inc., La Jolla, CA).

### Isothermal titration calorimetry (ITC)

For the ITC measurements, purified recombinant SPN-ARR fragment of SHANK3, GMPPCP-loaded His6- KRASQ61H and the RBD domain of human BRAF were exchanged into the ITC buffer containing 20 mM Tris pH 7.5, 500 mM NaCl, 0.5 mM TCEP (tris-carboxyethyl-phosphine) and 5 mM MgCl2. ITC experiments were performed using an ITC-200 (Microcal). Protein concentrations were estimated from UV absorbance at 280 nm. ITC titrations were performed at 25°C using 20 µM of SHANK3 with 350 µM of KRASQ61H, and 20 µM of KRASQ61H with 200 µM of RAF-RBD. Data were integrated and fitted to a single-site binding equation using Origin 7 software with ITC module (Microcal).

### Atomistic simulation models and methods

The simulated systems and their key details are described in the Extended Data Methods and in Extended Data Table 1. Briefly, SHANK3 was simulated with three different lipid bilayer systems to probe spontaneous membrane binding capabilities of SHANK3. Spontaneously formed KRAS-membrane complexes were also analysed. Finally, the KRAS-SHANK3 complex was obtained by aligning the structures of SHANK3 SPN (*30*) to the RAF RBD (*36*) coordinates. To run the simulations, we used the GROMACS simulation package version 2020 (*55*). Initiation of the systems followed the general CHARMM-GUI protocol (*56*). The simulation systems were first energy-minimised and then equilibrated with position restraints acting on the solute atoms.

### Immunofluorescence

To study co-localization of SHANK and mutant KRAS, cells were plated on glass-bottom dishes (Cat. no. P35G- 1.5-20-C, MatTek corporation), previously coated with fibronectin (5µg/ml) and collagen (10 µg/ml) overnight at 4°C, and transiently plasmid transfected using Lipofectamine® 3000 (Cat. no. L3000-015, Invitrogen) for 24h. U2OS cells were transfected with mRFP-SHANK3 wild-type (WT) and GFP-KRASG12V. Cells were then fixed with 4% PFA in phosphate buffered saline (PBS) for 10 min at RT and washed with PBS. Imaging was performed with a 3i spinning disk confocal (Marianas spinning disk imaging system with a Yokogawa CSU-W1 scanning unit on an inverted Carl Zeiss Axio Observer Z1 microscope, Intelligent Imaging Innovations, Inc.).

To study subcellular localization of ERK after SHANK3-KRAS interaction, MIA PaCa-2 cells were grown on coverslips overnight and transfected with GFP-tagged SPN WT or SPN R12E/K22D using Lipofectamine® 3000 (Invitrogen) for 24 h as described above. Cells were then fixed with 4% PFA in PBS for 10 min at RT, washed with PBS and permeabilised with 0.5% Triton-X-100 in PBS for 10 min at RT. PFA was quenched by incubating with 1 M Glycine for 30 min at RT. Cells were stained with the primary antibody diluted in PBS (ERK1/2, Cell Signaling, 91025, 1:100) for 30 min at RT. Cells were then washed and incubated with Alexa Fluor-conjugated secondary antibody (1:300, Life Technologies) and 4′6-diamidino-2-phenylindole (DAPI, nuclei staining, 1:10000; Cat. no. D1306, Life Technologies) diluted in PBS for 30 min at RT. Finally, cells were washed, and imaging was performed with a 3i spinning disk confocal (Marianas spinning disk imaging system with a Yokogawa CSU-W1 scanning unit on an inverted Carl Zeiss Axio Observer Z1 microscope, Intelligent Imaging Innovations, Inc.). Samples were either imaged right away or stored at +4°C in the dark until imaging. In order to obtain a quantitative estimate for the extent of ERK nuclear translocation (indicative of ERK activity), captured images of cells were then analysed by calculating the ratio of staining intensity measured in the nucleus to that of a cytoplasmic region of the cell. This procedure takes into account the potential variability in staining efficiency between different cell cultures.

To use the ERK biosensor in imaging of ERK activity, we relied on the ERK-KTRmRuby2 probe (*41*). For lentiviral transduction of the ERKKTR into PANC1 cells, HEK293T packaging cells were co-transfected with pMDLg/pRRE (Plasmid #12251, Addgene), pRSV-Rev (Plasmid #12253, Addgene), pMD2.G (Plasmid #12259, Addgene) and pLentiPGK-Blast-DEST-ERKKTRmRuby2 (Plasmid # 90231, Addgene; a kind gift from Markus Cover (*41*), using Lipofectamine 3000 as per the manufacturer’s instructions. Packaged lentiviruses were then applied to PANC-1 cells in the presence of polybrene (8 µg/ml, TR-1003-G, Sigma-Aldrich) and incubated overnight. Successfully transduced cells were then selected using blasticidin (6 µg/ml, Cat. no. 15205, Sigma-Aldrich). For the experiments, PANC-1 ERKKTR cells were plated on glass-bottom dishes (Cat. no. P35G-1.5-20- C, MatTek Corporation) and silenced by siRNAs for two or three days, as described above. Then, cells were fixed with 4% PFA in PBS for 10 min at RT and washed with PBS. Imaging of ERK-KTR was performed with a 3i spinning disk confocal (Marianas spinning disk imaging system with a Yokogawa CSU-W1 scanning unit on an inverted Carl Zeiss Axio Observer Z1 microscope, Intelligent Imaging Innovations, Inc.). Samples were either imaged right away or stored at +4°C in the dark until imaging. Analysis of cytoplasmic to nuclear (C/N) ratios of the ERKKTR reporter was performed in ImageJ (National Institutes of Health) by measuring the integrated density of fluorescence in the representative area of cytoplasm versus the representative area of nucleus.

### FRET imaging using fluorescence lifetime imaging microscopy (FLIM)

To visualise SHANK3-KRAS interaction in cells, we used FRET-FLIM. HEK293 cells were grown on coverslips overnight and transfected with an mGFP-tagged donor construct (GFP-SHANK3 SPN WT or GFP SHANK3 SPN R12E/K22D) and mCherry-tagged acceptor construct (mCherry-KRASG12V) using Lipofectamine® 3000 (Invitrogen) or jet PRIME (Polyplus) transfection reagent. Cells were fixed with 4% PFA/PBS and mounted with Mowiol 4-88 on microscope slides 24-48 h post-transfection. Fluorescence lifetimes of the GFP-tagged donor constructs were measured using a fluorescence lifetime imaging attachment (Lambert Instruments, Leutingwolde, The Netherlands) on an inverted microscope (Zeiss Axio Observer.D1). Fluorescein (0.01 mM, pH 9) was used as a lifetime reference standard. In addition, it served to calibrate a fixed setting that allows acquisition of data from cells with comparable expression levels. Three independent biological repeats were performed, and the apparent fluorescence resonance energy transfer (FRET) efficiency was calculated from obtained fluorescence lifetimes. The percentage of the apparent FRET efficiency (Eapp) was calculated using the measured lifetimes of each donor-acceptor pair (τDA) and the average lifetime of the donor only (τD) samples (Eapp=(1 − τDA/τD) × 100%) (*37*).

### Effector-recruitment FRET assay

HEK293 cells were first silenced with control or *SHANK3* targeting siRNA for 48 hours, and then, seeded on a 6-well plate with glass coverslips, and plasmid-transfected with the donor alone (mGFP-tagged KRASG12V construct) in control samples, or together with the acceptor mRFP-RBD in CRAF-RBD-recruitment FRET experiments. After 48 h of plasmid transfection, coverslips were fixed with 4% PFA/PBS for 15 min and then washed with PBS, and coverslips were mounted with Mowiol 4–88 (Sigma-Aldrich) on microscope slides. The mGFP fluorescence lifetime was measured using a fluorescence lifetime imaging attachment (Lambert Instruments) on an inverted microscope (Zeiss AXIO Ovserver.D1) as previously described in (*37*).

### Cell viability assays

AnnexinV-FITC/PI flow cytometry assay and annexinV-FITC imaging-based assay were used to evaluate apoptotic and necrotic cell death in cells cultured in monolayers and as spheroids. One to three days after silencing, cells were stained by Annexin V-FITC Apoptosis Detection Kit (Cat. no. BMS500FI-100, eBioscience™) according to the manufacturer’s instructions. Apoptotic cells were detected using BD LSR Fortessa™ analyser (BD Biosciences). The effects of anti-SHANK3 SPN nanobody (see below) expression on cell viability were tested in three *KRAS*-mutant cell lines that we have established to be sensitive to *SHANK3* silencing induced cell death (PANC-1, A549, H441). mCherry-tagged anti-SHANK3 nanobodies (A01 and E01) or control-mCherry were transiently expressed in the cells lines, and apoptotic cell fraction was assessed with Annexin-V-FITC labelling (without PI staining) and flow cytometry four days post-transfection.

Cell viability of *SHANK3*-silenced PANC-1 cells treated with ERK and MEK inhibitors were measured using Cell Counting Kit-8 (WST-8 / CCK-8; Cat. no. ab228554, Abcam). Cells were silenced for 24 h as described above and then seeded on a 96-well plate with full medium containing DMSO (control), Selumetinib (ADZ6244, Selleckchem) or a selective ERK1/2 inhibitor (SCH772984, Selleckchem) at different concentrations (concentrations used: 0, 0.1 nM, 0.5 nM, 1 nM, 5 nM, 10 nM, 31.3 nM, 62.5 nM, 124 nM. 250 nM, 500 nM and 1 µM). Cell viability was measured 96 h after silencing (drugs for 72 h).

3D spheroid formation was performed as described above. Cells were embedded between two layers of Matrigel and covered with full cell culture medium. The spheroid formation and growth was followed by IncuCyte imaging. SHANK3-depletion was induced by doxycycline (dox) in established sh*SHANK3* PANC-1 spheroids. Both doxycycline (+dox; 2 µg/ml) and AnnexinV (1:200, Annexin V-FITC Apoptosis Detection Kit, Cat. no. BMS500FI-300, eBioscience™) were added to established spheroids at day 5 and spheroid growth was followed for 10 days. Culture medium containing dox (or –dox) and AnnexinV was changed every second day. Analysis was performed using IncuCyte S3 software. Alternatively, the spheroid growth/viability was measured using Cell Counting Kit-8 (WST-8 / CCK-8; Cat. no. ab228554, Abcam). On day 15 of 3D spheroid culture (after 10 days of dox-induction), medium was aspirated and fresh medium supplemented with 1:10 of WST-8 solution. After two hours of incubation protected from light at 37°C, absorbance was measured at 460 nm.

### *In vivo* chick embryo chorioallantoic membrane (CAM) assay

Fertilised chicken eggs were incubated as previously described (*57*). Briefly, the eggs were washed, and the development was started by placing the eggs in a 37 °C incubator. On day 3 of development, a small hole was made in the eggshell to drop the CAM. On developmental day 7, a plastic ring was placed on the CAM and one million cells (transiently plasmid or siRNA transfected) were implanted inside the ring in 20 μl of 50% Matrigel (Cat. no. 354230, Corning) diluted in PBS. After 4-5 days, tumours were dissected, weighed, and fixed in 10% formalin.

### Generation of doxycycline-inducible sh*SHANK3* PANC-1 cell line

SMART lentiviral shRNA vectors for doxycycline-inducible suppression of human *SHANK3* gene expression were purchased from Dharmacon as viral particles (Dox-inducible SMARTvector sh*SHANK3*, V3SH7669-228381856, Dharmacon). Packaged lentiviruses (40 MOI) were applied to PANC-1 cells in the presence of polybrene (8 µg/ml, Cat. no. TR-1003-G, Sigma-Aldrich) for 48 h; first, cells were incubated with transduction mix in serum-free medium for 20 h, and then full medium was added to cells without removing the transduction mix. Two days after transduction, the medium was replaced with full medium and cells were cultured for additional 48 h. Four days after transduction cells were selected using puromycin (4 µg/ml, Cat. no. 15205, Sigma-Aldrich). Single-cell clones were created by screening for high induction efficacy (bright tRFP positive clones after dox-induction; indicative of *SHANK3* shRNA expression). All established PANC-1 sh*SHANK3* expressing cell lines (single and a mix of clones 1C and 4S) were cultured in DMEM supplemented with 10 % FBS and 2mM L-glutamine, 2 µg/ml puromycin. Cells were regularly tested and found to be free from mycoplasma contamination. For experiments, doxycycline induction (+Dox; 1-2 µg/ml) was started 24 hours post-plating. Culture medium including doxycycline (+Dox) (or –Dox) was changed every second day.

### Subcutaneous tumour xenografts in Nude mice

To evaluate the requirement for SHANK3 in established tumours, six- to eight-week-old female athymic Nude mice (Hsd:Athymic Nude-foxn1nu, Envigo, France) were injected in the flank with 5 million PANC-1 doxycycline-inducible *SHANK3* shRNA-containing cells (pool of clones 4S and 1C) resuspended in 100 µl PBS with 50% Matrigel (Cat. no. 354230, Corning). When tumours reached an average mean volume of 100 mm^3^, the mice with similarly sized tumours were blindly randomised into cohorts. Then, mice were fed normal chow (control group; Teklad 2914 diet, Envigo) or doxycycline-containing chow (*SHANK3* depletion induced; Teklad doxycycline-diet, 625 mg/kg, in 2014 diet base, irradiated (2914), color red, Envigo) daily. In addition, mice received two intraperitoneal (i.p) injections of PBS or doxycycline (80 mg/kg of body weight). Successful induction of *SHANK3* shRNA expression was confirmed by IVIS imaging (tRFP expression after dox-induction; indicative of *SHANK3* shRNA expression). Tumours were measured with digital calipers twice a week and tumour volumes were calculated according to the formula V = (π/6)(d1 × d2)^3/2, where d1 and d2 are perpendicular tumour diameters. Mice were sacrificed at day 26 days post-induction (74 post-engraftment), and tumours were dissected, weighed and snap frozen in liquid nitrogen for mRNA isolation.

All animal experiments were ethically assessed and authorised by the National Animal Experiment Board and in accordance with The Finnish Act on Animal Experimentation (Animal license numbers ESAVI/9339/2016 and ESAVI/37571/2019).

### Immunohistochemistry analysis of tumours

Formalin-fixed, paraffin-embedded tissue samples were cut to 4 µm sections, deparaffinised and rehydrated with standard procedures. For immunohistochemistry (IHC) of CAM tumours, heat-mediated antigen retrieval was done in citrate buffer (pH 6 for cleaved caspase-3, pH 9 for Ki-67). Sections were washed with washing buffer (0.05 M Tris-HCl pH 7.6, 0.05 % Tween20), blocked for endogenous hydrogen peroxide activity, and incubated with Normal Antibody Diluent (NABD; Cat. No. BD09-125, Immunologic). Sections were then incubated with a Ki-67 antibody (Cat. no. AB9260, Millipore, diluted 1:1000) or a Cleaved Caspase-3 (Asp175) antibody (Cat. no. 9664, clone 5A1E, Cell Signaling Technology, diluted 1:500) for 1 h. After washes, samples were incubated 30 min with a BrightVision Goat anti-Rabbit HRP (Cat. no. DPVR110HRP, Immunologic) secondary antibody, and washed again. After washes, DAB solution (Cat. no. K3468, DAKO) was added for 10 sec followed by washing. After counterstain with Mayer’s HTX, slides were dehydrated, cleared in xylene and mounted in Pertex. Stained samples were imaged with Pannoramic P1000 Slide Scanner (3DHISTECH Ltd) and analysed with QuantCenter software with NuclearQuant quantification module (3DHISTECH Ltd).

### Recombinant nanobodies against SHANK3

Nanobodies (single domain antibodies) against SPN were generated by Hybrigenics Services SAS (Evry, France; www.hybribody.com) by three rounds of Phage Display selection of their naïve VHH-library against recombinant biotinylated GST-SPN protein. Chosen nanobodies in the pHEN2 vector were then produced as recombinant proteins in BL21 bacteria using IPTG induction and purified according to Hybrigenics’ protocol. Details of nanobody selection and purification are described in the Extended Data Methods.

### Functional testing of anti-SHANK3 nanobodies

The ability of the anti-SHANK3 SPN nanobodies (A01 and E01) to disrupt SHANK3 SPN-KRAS-GTP interaction was tested using two independent assays, as detailed in the Extended Data Methods. Briefly, in the ELISA system purified recombinant His-SPN SHANK3 protein was used to coat wells and then preincubated with the anti-SHANK3 or irrelevant control nanobodies before addition of GST-tagged purified recombinant KRAS protein loaded with non-hydrolyzable GTP analogue. An Europium (Eu)-labeled anti-GST antibody was then used to detect the bound KRAS protein using time resolved fluorescence plate reader.

In the pull-down assay His-SHANK3 SPN protein was bound to Ni-resin before incubating with the SHANK3 SPN nanobodies. Thereafter, GST-KRAS-GTP or GST alone was incubated with the beads under rotation at 4 ℃. After washings, the eluted GST-proteins were detected using immunoblotting with anti-GST antibodies.

### Statistical analyses

The sample size for studies was chosen according to previous studies in the same area of research. The GraphPad program was used for all statistical analyses. Normal distribution of the data was tested with D’Agostino & Pearson omnibus normality test. Student’s t-test (unpaired, two-tailed) with Welch’s correction was used for two groups when normality could be confirmed. Nonparametric Mann–Whitney U-test was used when two non-normally distributed groups were compared or when normality could not be tested [due to a too small data set (n < 8)]. ANOVA with Holm-Sidak’s or Dunnett’s multiple comparison test was used when comparing more than two normally distributed groups. Kruskal-Wallis non-parametric test with Dunn’s multiple comparison test was used when comparing more than two non-normally distributed groups. Data are presented in column graphs or scatter dot plots with mean ± standard error of mean (s.e.m) or mean ± standard deviation (s.d) and P-values. Individual data points per condition are shown and n-numbers are indicated in figure legends. Superplots show data of all measurements and the average of each biological replicate. P-values less than 0.05 were considered to be statistically significant.

## Data and materials availability

The authors declare that the data supporting the findings of this study are available within the paper and its supplementary information files.

## Acknowledgements

We thank P. Laasola, J. Siivonen, E-M. Vesilahti, M. Miihkinen, S. Salomaa and A. Isomursu for technical assistance and scientific discussion, the Ivaska lab for critical reading and feedback on the manuscript and O. Pentikäinen for protein complex modelling. The Cell Imaging and Cytometry Core (Turku Bioscience Centre, University of Turku) and Turku Centre for Disease Modelling (TCDM), both supported by Biocenter Finland, the Euro-BioImaging Finnish Node (Turku Finland), the University of Turku Histocore and Genome Editing core are acknowledged for services, instrumentation, and expertise. This study has been supported by the Academy of Finland (325464 J.I., G.J. and E.P.), the Academy of Finland CoE for Translational Cancer Biology (J.I.), an ERC CoG grant (no. 615258; J.I.), the Sigrid Juselius Foundation (J.I. and G.J., and E.P.), the Finnish Cultural Foundation (J.L. and E.P) and the Finnish Cancer Organization (J.I., M.S., GJ). J.L and J.K have been supported by the Turku Doctoral Programme of Molecular Medicine (TuDMM), J.L by the Instrumentarium Foundation, the Orion Research Foundation sr and the K. Albin Johanssons Foundation, and J.R.W.C. is supported by the European Union’s Horizon 2020 research and innovation programme under the Marie Sklodowska-Curie grant agreement [841973]. AKN was supported by CRUK Beatson Institute core funding A17196, A31287 - awarded to OJS. OJS was supported by CRUK grants A21139, A12481, A17196 and A31287 and ERC Starting grant 311301. The I.V. group is supported by the Sigrid Juselius Foundation, the Academy of Finland (project no. 331349), Human Frontier Science Program (project no. RGP0059/2019), the Helsinki Institute of Life Science (HiLIFE) Fellow program, and Cancer Foundation Finland. P.R. is supported by Drug Research Doctoral Programme at University of Turku. We also gratefully acknowledge CSC – IT Center for Science (Espoo, Finland) for providing ample computing resources.

## Author contributions

Conceptualization: JL, JI. Methodology: JL, JRWC, UP, IB, JV, DA, IV, EP, JI. Formal Analysis: JL, JK, UP, IB, JV, EP, HP. Investigation, JL, JK, HP, TV, GJ, EP, JV, PR, IMDP, EW, AKN. Visualization: JL, HH. Writing – Original Draft: JL, MS, JI. Writing – review & editing: JL, HH, JRWC, GJ, EP, AKN, OJS, DA, JV, IV, MS, JI. Supervision: OJS, IB, IV, DA, UP, JI. Funding Acquisition: IV, JI.

## Extended Data information

**Extended Data Figure 1.**
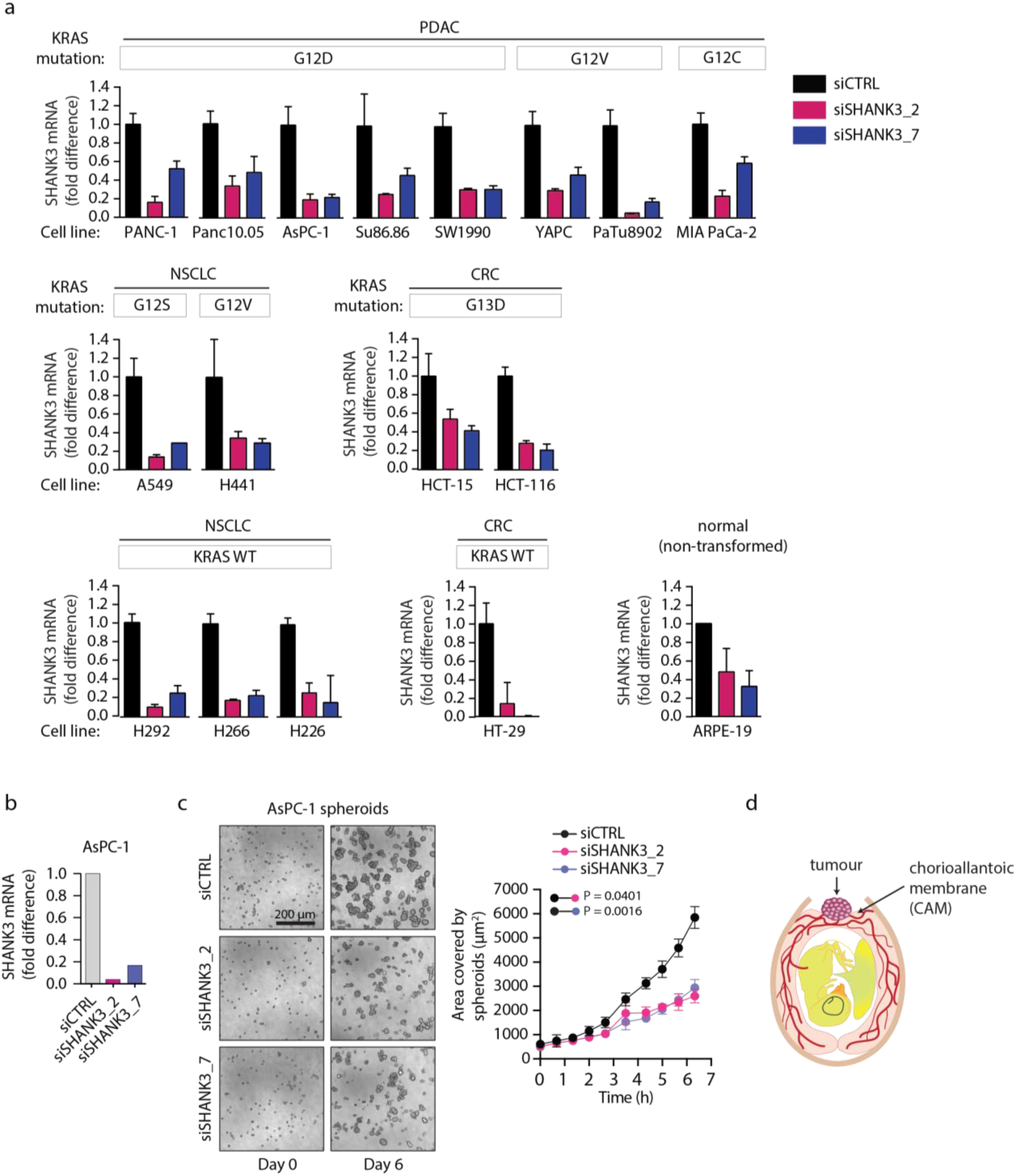
*SHANK3* depletion impairs KRAS-driven cancer cell proliferation and tumour growth, Related to Figure 1. **a,** *SHANK3* expression (mRNA levels; the mean ± s.d) of control (siCTRL) and *SHANK3*-silenced (siSHANK3_2 and siSHANK3_7 are two different siRNAs) cancer cell lines with distinct *KRAS* mutations (PDAC: PANC-1, Panc10.05, AsPC-1, Su86.86, SW1990, YAPC, PaTu8602 and MIA PaCa-2; NSCLC: A549 and H441; CRC: HCT-15 and HCT-116) or with wild-type *KRAS* (NSCLC: H292, H226 and H226; CRC: HT-29) and non-transformed cells (ARPE-19). b,c, Spheroid growth of control or *SHANK3*-silenced *KRAS*-mutant AsPC-1 cells. b, *SHANK3* mRNA levels. c, Representative images and quantification (mean ± s.d.) of the spheroid areas from n = 3 independent experiments. Unpaired Student’s t-test with Welch’s correction (at the endpoint). d, A schematic illustration of the chick embryo chorioallantoic membrane (CAM) xenograft assay.

**Extended Data Figure 2.**
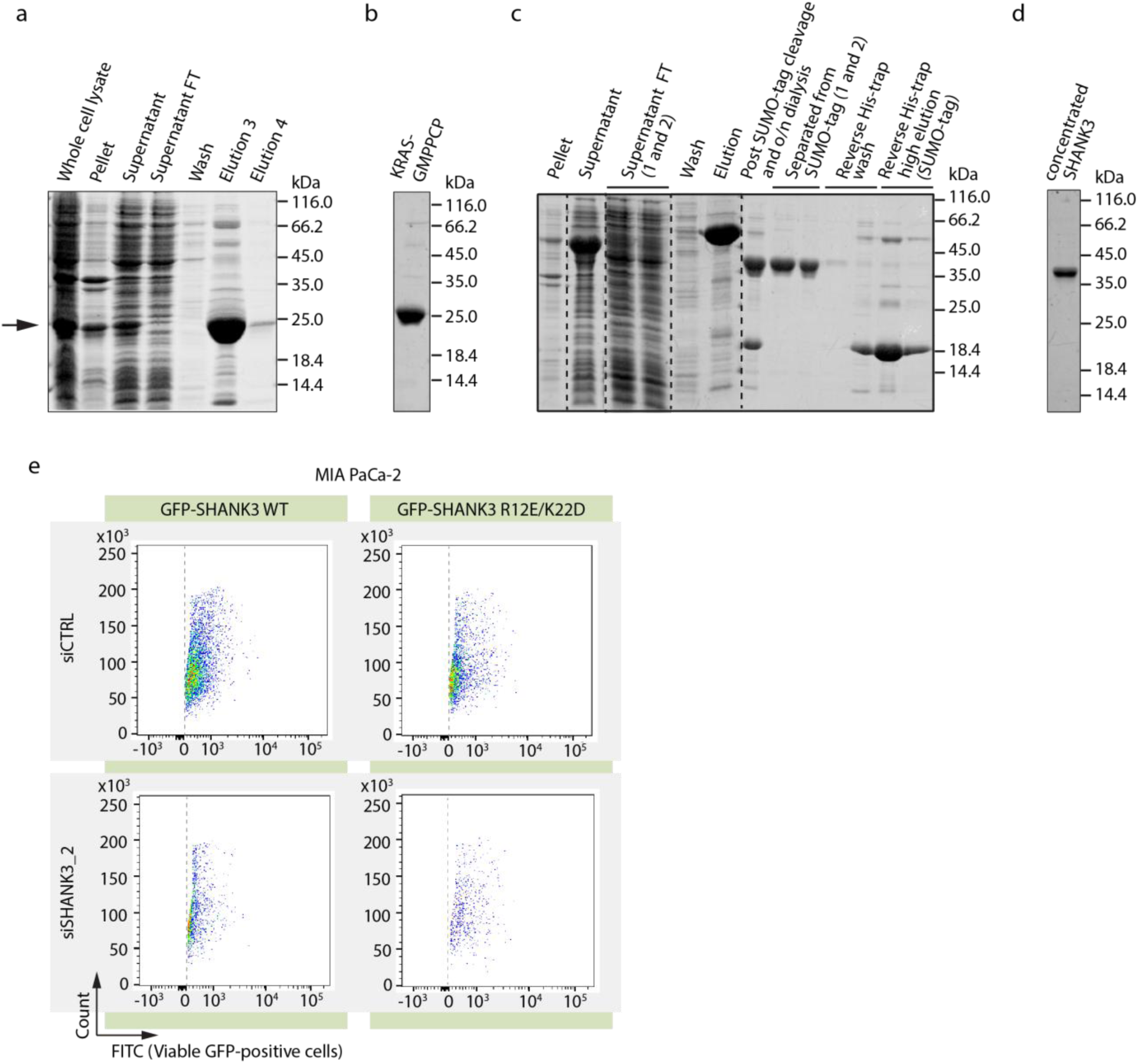
SHANK3 interacts directly with active/mutant KRAS, Related to Figure 2. **a,** KRASQ61H purification by Ni^2+^-NTA affinity chromatography. Supernatant FT = supernatant flow-through from His-trap column. Arrow points to the expected mw of KRASQ61H. b, Pure KRASQ61H after overnight exchange with GMPPCP used for ITC. c, SHANK3 WT 1-348 purification. 6xHis-SUMO-tag removed via overnight dialysis and cleavage with SUMO protease, followed by a second Ni^2+^-NTA affinity chromatography to separate the tag from the protein. d, Concentrated sample of SHANK3 WT 1-348 used for ITC. e, Representative scatter plots of viable GFP positive MIA PaCa-2 cells expressing GFP-tagged wild-type or RAS-binding deficient (R12E/K22D) SHANK3 after silencing of endogenous human *SHANK3* (36 h).

**Extended Data Figure 3.**
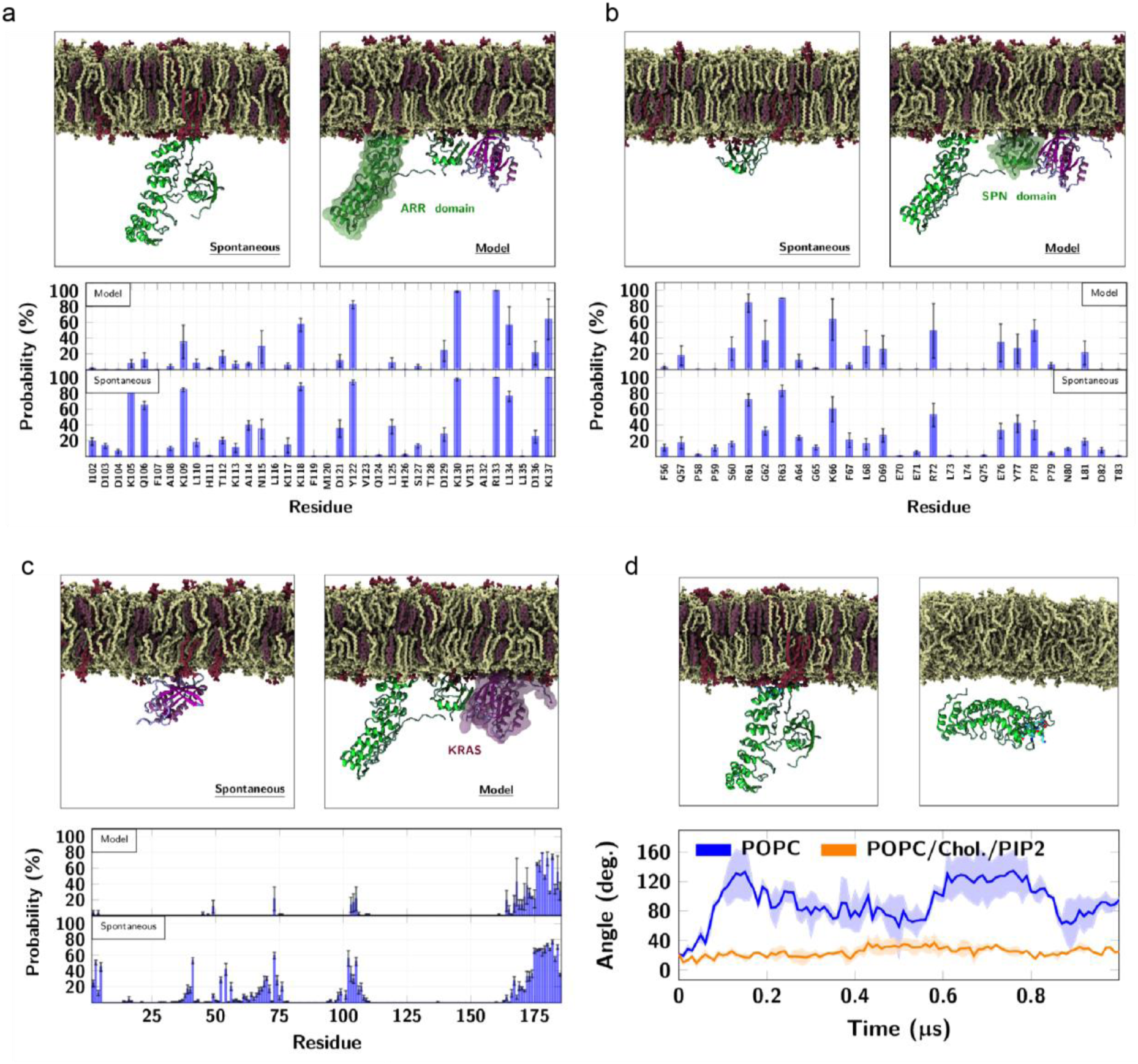
SHANK3 competes with RAF for the active KRAS binding, Related to Figure 3. **a,b,** SHANK3 SPN-ARR interaction with lipid bilayers. a, Final simulation frames (1000 ns) resulting from spontaneous membrane binding of isolated SHANK3 SPN-ARR (left, System S2) and modelled membrane binding of SHANK3 SPN-ARR with KRAS (right, System S5). Bottom image shows the probability of contacts (minimum distance < 0.6 nm) between the residues of SHANK3 and PIP2 lipids and indicates which residues of the ARR domain contact the lipids in the two cases. Only residues at the membrane binding region are listed. b, Final simulation frames (1000 ns) resulting from spontaneous membrane binding of isolated SHANK3 SPN (left, System S3) and modelled membrane binding of SHANK3 SPN-ARR with KRAS (right, System S5). Bottom image shows the probability of contacts (minimum distance < 0.6 nm) between the residues of SHANK3 and PIP2 lipids, indicating which residues of the SPN domain contact the lipids in the two cases. Only residues at the membrane binding region are listed. Errors are standard errors of mean. c, The binding of KRAS on lipid membranes observed in the simulations. Final simulation frame (1000 ns) from a simulation of KRAS with a PIP2- containing bilayer (left, System S4), indicating its spontaneous binding, and final frame (1000 ns) of modelled membrane binding of SHANK3 SPN-ARR with KRAS (right, System S5). Bottom image shows the probability of contacts (minimum distance < 0.6 nm) between the residues of KRAS and PIP2 lipids, indicating which residues of KRAS contact the lipids in the two cases. Errors are standard errors of mean. d, Final simulation frame (1000 ns) from a simulation of SPN-ARR with a PIP2-containing bilayer (left, System S2) and the final frame (1000 ns) from a simulation of SPN-ARR with a POPC (1-Palmitoyl-2-oleoyl-sn-glycero-3-phosphocholine)/ Phosphatidylinositol 4,5-bisphosphate/ Cholesterol) bilayer (right, System S1). Bottom image shows results for the tilt angle of the vector between the Cα atoms of the SHANK3 residues 114 and 286. These residues were chosen to span the long axis of the ARR domain (long axis of the protein). Zero degrees corresponds to the alignment of the vector with the bilayer normal. The data were calculated with the *gmx bundle* tool within the GROMACS package.

**Extended Data Figure 4.**
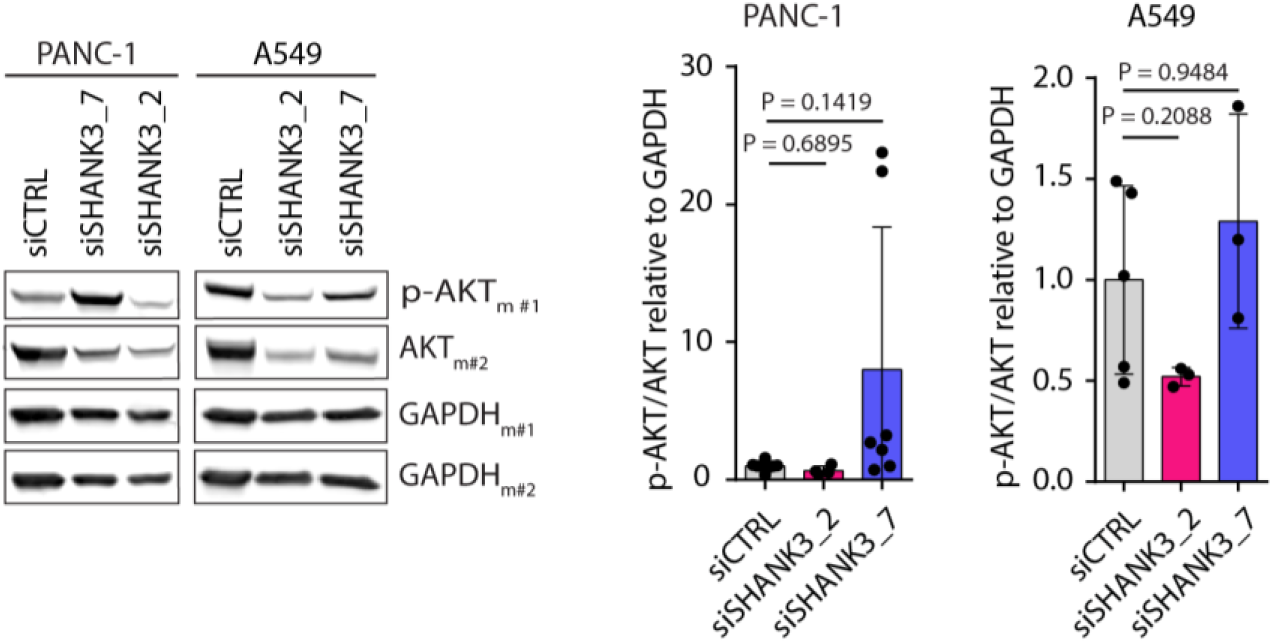
Loss of SHANK3 has no significant effects on AKT signalling in *KRAS*-mutant cells, Related to Figure 4. Analyses of AKT signalling in *SHANK3*-silenced *KRAS*-mutant in PANC-1 and A549 cells. Left: Representative immunoblots from PANC-1 and A549 cells analysed three days after silencing. AKT, total AKT; p-AKT, phospho-AKT S473; GAPDH, a loading control (duplicate membranes: m#1 and m#2). Loading controls shown are from Fig. 4a. The efficiency of *SHANK3* silencing is shown in Fig. 5A. Right: Quantifications showing the levels of AKT phosphorylation (relative to total AKT). Data are the individual experiments and the mean±s.d. Kruskal-Wallis test and Dunn’s post hoc test.

**Extended Data Figure 5.**
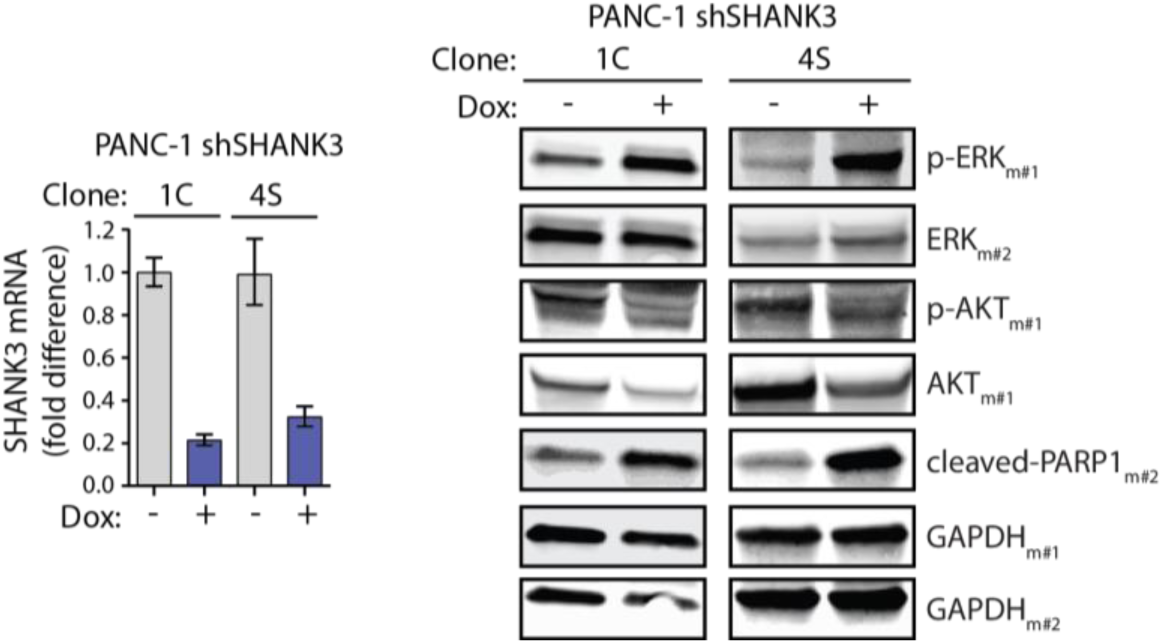
Inducible silencing of *SHANK3* activates ERK and induces apoptosis, Related to Figure 5. Left: *SHANK*3 gene expression (mRNA levels) showing the efficiency of *SHANK3* silencing in control or doxycycline-induced (Dox: +; 72 h) shSHANK3 expressing PANC-1 clones (clones 1C and 4S). Right: Representative immunoblots of the indicated proteins in control or doxycycline-induced sh*SHANK3* expressing PANC-1 clones collected three days after induction. Samples were resolved and blotted on duplicate membranes (m#1 and m#2). p-ERK, phospho-ERK1/2 (Thr202/Y204); ERK, total ERK; AKT, total AKT; p-AKT, phospho-AKT S473; cleaved-PARP1, indicative of apoptosis; GAPDH, a loading control.

**Extended Data Figure 6.**
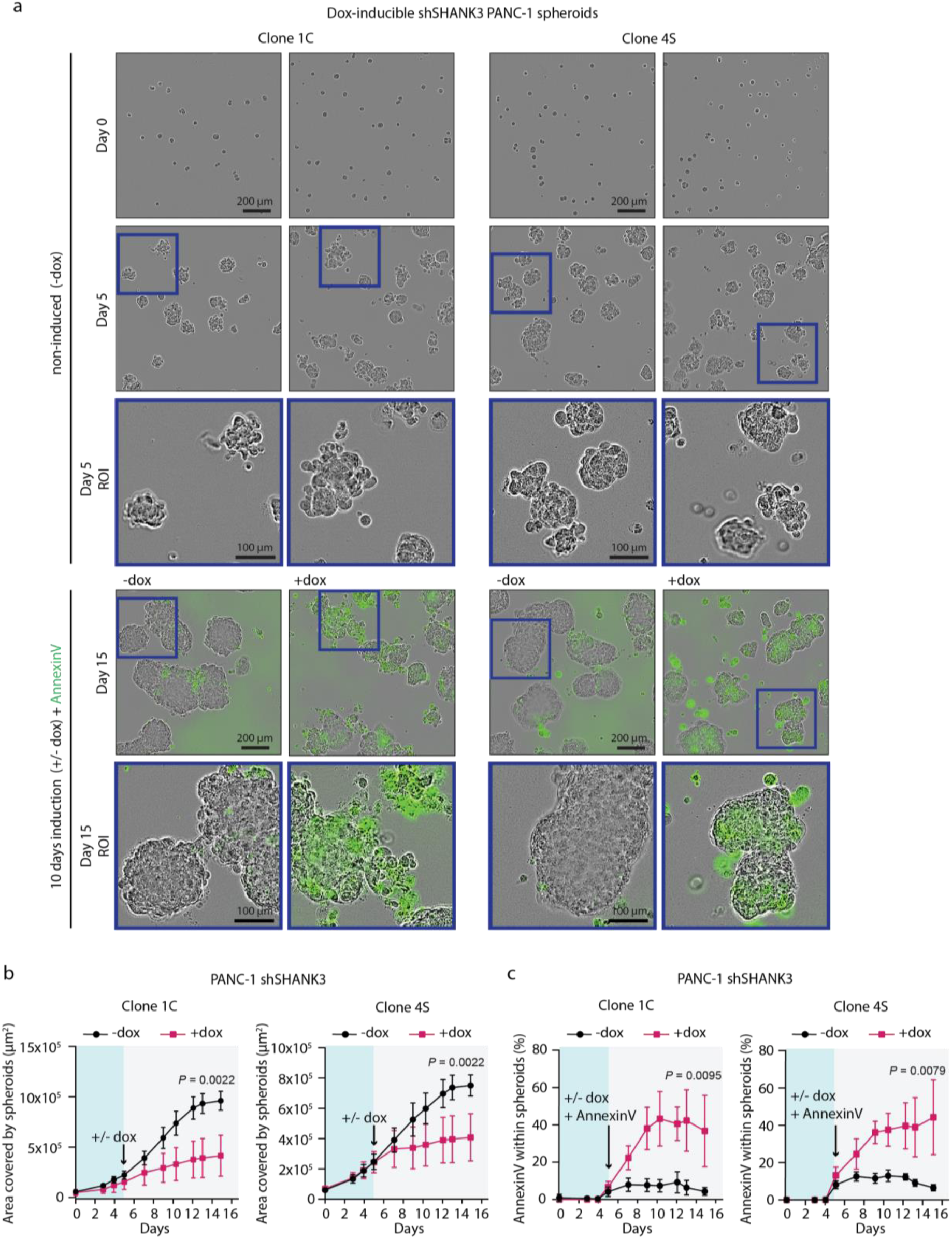
SHANK3 depletion in established *KRAS*-mutant spheroids induces apoptosis, Related to Figure 5. a, Representative images of spheroids and apoptosis (green AnnexinV positive cells) of control or doxycycline-induced (+dox) sh*SHANK3*-expressing PANC-1 clones (clones 1C and 4S); doxycycline-induction was started after 5 days of spheroid growth and continued until day 15. ROI, region of interest (within blue squares). b, Quantification of spheroid growth of control or dox-induced sh*SHANK3* expressing PANC-1 clones (clone 1C and 4S). Data represent mean ± s.d.; n = 6 measurements from 2 independent experiments. Mann-Whitney test. c, Analysis of apoptosis (AnnexinV positive area) in established control (-dox) or SHANK3-depleted (+dox) PANC-1 spheroids (clones 1C and 4S). Both doxycycline and AnnexinV were added to spheroids at day 5 (arrow). Data represent mean ± s.d.; n = 6 measurements from 2 independent experiments. Mann-Whitney test.

**Extended Data Figure 7.**
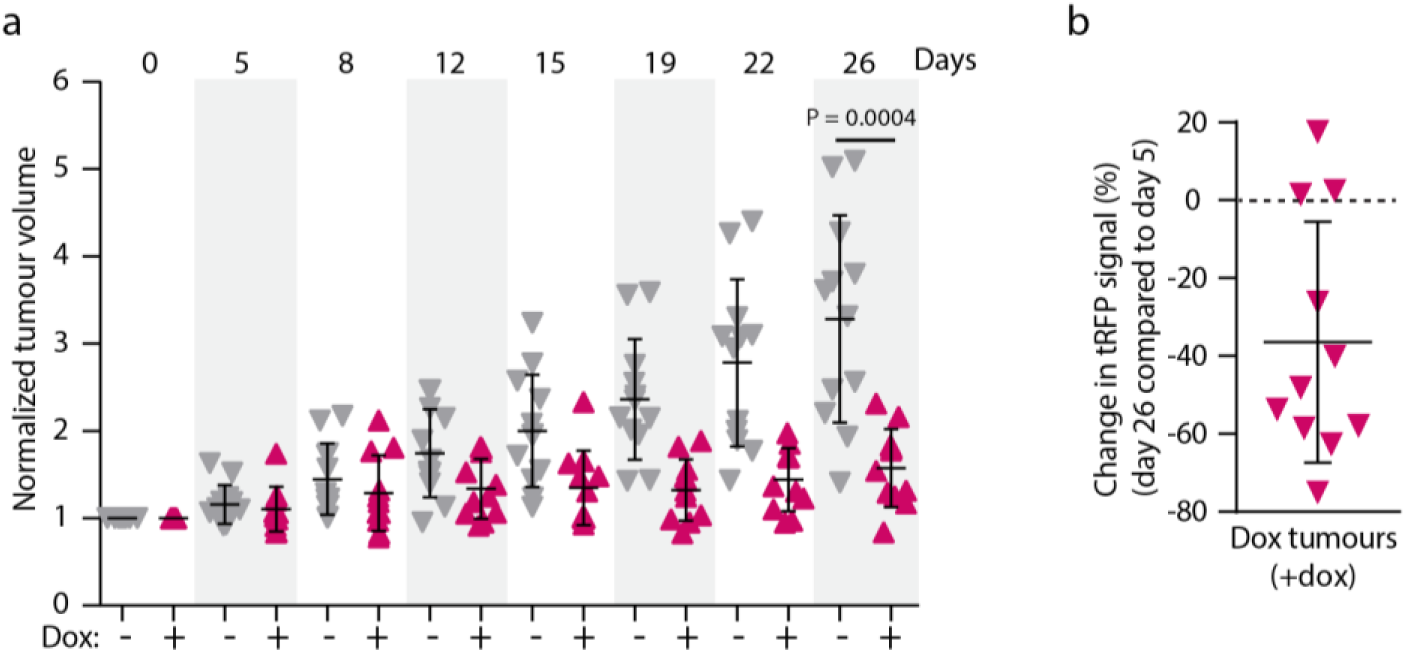
SHANK3 depletion in established PDAC tumours impairs tumourigenic growth *in vivo*, Related to Figure 5. **a,** Growth rate of subcutaneously injected PANC-1 cell xenografts with doxycycline-inducible SHANK3 knockdown (+dox) over the indicated time. Tumour growth was monitored with bi-weekly palpations. Data represent mean ± s.d. of n = 11 (+dox) and 12 (-dox) tumours. Unpaired Student’s t-test with Welch’s correction. b, SHANK3 depletion was observed by tRFP reporter (visual tracking of transduction and expression of shRNA). Shown is a change in tRFP signal (%) in tumours with doxycycline-inducible SHANK3 knockdown (Dox) at the end of the experiment. Data represent mean ± s.d. of n = 11 (+ dox). Each data point represents an individual mouse (a,b).

**Extended Data Table 1.** Key details of the simulated systems, Related to Figure 3.

| System | Protein components<br>(mutation, residue<br>range, other information) | Number of lipids<br>(POPC/ cholesterol/<br>PIP2) | Number of water<br>molecules, and<br>number of ions<br>(K<sup>+</sup>,Cl<sup>-</sup>) | Number of<br>replicas x<br>duration (ns) | PDB Ref. |
| --- | --- | --- | --- | --- | --- |
| **S1** | SHANK3 SPN-ARR (wt,<br>2–347) | 500/0/0 | 63047<br>(173, 171) | 4 x 1000 | 5G4X |
| **S2** | SHANK3 SPN-ARR (wt,<br>2–347) | 364/168/28 | 61451<br>(282, 168) | 4 x 1000 | 5G4X |
| **S3** | SHANK3 SPN<br>(wt, 2–93) | 182/84/14 | 24276<br>(121, 66) | 4 x 1000 | 5G4X |
| **S4** | KRAS<br>(wt, 1–185) | 166/76/14 | 29032<br>(132, 78) | 4 x 2000 | 6PTW |
| **S5** | SHANK3 SPN-ARR (wt,<br>5–363)<br>KRAS (wt, 1–185) GNP | 414/190/26 | 74534<br>(303,204) | 4 x 1000 | 6PTW /<br>6KYK |

## Extended Data Methods

### Production and purification of recombinant proteins

#### KRASG12V

The sequence of the synthetic gene of *KRAS* 4B was designed according to *E.coli* codon usage. *KRAS* was PCR amplified and cloned to a modified pGEX vector (GE Healthcare, Chicago, IL). The G12V mutation to KRAS 4B was ordered from BioCat (https://www1.biocat.com).

Glutathione S-transferase (GST) fusion KRASG12V protein was expressed in Terrific Broth (2.4 % w/v yeast extract, 1.2 % w/v tryptone, 0.5 % w/v glycerol, 0.017 M KH2PO4, 0.072 M K2HPO4, 100 µg/ml Ampicillin) by the addition of isopropyl-β-D-1-thiogalactopyranoside to 0.4 mM at 20℃ for 20 h using E. Coli BL21 Gold cells. The cells were lysed by sonication on ice (Sonopolus 4000) at 40 % amplitude (4x, 1 second pulse on and 1s pulse off) for 1 minute and subsequently centrifuged at 35,000 g for 30minutes at 4 ℃ to clear the lysate. The GST KRASG12V fusion protein was purified with Protino Glutathione Agarose 4B (Macherey-Nagel, Düren, Germany) and GST was cleaved by Tobacco Etch Virus (TEV) protease (Invitrogen, Life Technologies, Carlsbad, CA) at 4 °C for 16 hours. The TEV protease cleavage extended KRASG12V construct in the N-terminal by four amino acid residues, G, A, M and G. The proteins were further purified by size exclusion chromatography (SEC) with a HiLoad 26/60 Superdex 200 pg column (GE Healthcare, Chicago, IL) in SEC buffer (50 mM Tris, pH 7.3, 300 mM NaCl, 1 mM DTT, 0.1 % CHAPS) using an ÄKTA pure chromatography system (GE Healthcare). The protein was concentrated with Amicon ultracentrifugal 10K filter device (Millipore, Sigma, Burlington, MA). The homodispersity of the proteins was verified with sodium dodecyl sulphate polyacrylamide gel electrophoresis (SDS-PAGE).

#### KRASQ61H

The plasmid containing human His6-KRASQ61H activating oncogenic mutant (residues 1-169) was a gift from Cheryl Arrowsmith (Plasmid #25153, Addgene). The protein was expressed using *E.coli* BL21(DE3) competent cells (Invitrogen) cultured in Luria Broth (LB) medium. Cells were grown at 37°C in LB medium supplemented with antibiotics to an OD600 of 0.6, cooled to 18°C and induced using 300 μM IPTG (Isopropyl β-D-1-thiogalactopyranoside) for 16 hours. Cells were pelleted by centrifugation and resuspended in 20 mM Na2HPO4 pH 7.4, 500 mM NaCl, 5 mM MgCl2, 2 mM DTT and 25 mM imidazole, treated with protease cocktail inhibitor VII (Cat. no. 539138, Calbiochem) and sonicated on ice. The protein was purified using nickel-affinity chromatography with a linear gradient of lysis buffer containing 500 mM imidazole, but without DTT. Immediately after purification, 2 mM DTT was added to the protein fractions. Protein purity was checked by SDS-PAGE gel. Bound nucleotide was exchanged for non-hydrolysable GTP analogue GMPPCP using alkaline phosphatase beads (Sigma-Aldrich) and following the protocol of John et.al. (1).

### RBD domain of B-Raf

The RBD domain of human BRAF (Uniprot P15056) corresponding to residues Ser151-Leu232 was cloned into pOPNB vector (OPPF-UK) from codon-optimised synthetic DNA at GeneMill facility, University of Liverpool. The protein was expressed using BL21 competent cells (Invitrogen) cultured in LB and purified using Ni-NTA column with standard protocol. His-tag was cleaved with recombinant His-tagged 3C protease and removed by a reverse pass on the Ni-NTA column.

#### SPN domain of SHANK3

His-tagged SPN protein was expressed in Terrific Broth (2.4 % w/v yeast extract, 1.2 % w/v tryptone, 0.5 % w/v glycerol, 0.017 M KH2PO4, 0.072 M K2HPO4, 100 µg/ml Ampicillin) by the addition of isopropyl-β-D-1-thiogalactopyranoside to 0.4 mM at 18℃ for 20 h using *E. coli* BL21 Gold cells. Prior to the cell lysis, small amounts of both lysozyme and DNase were added and then the cells were lysed by sonication on ice (Sonopolus 4000) at 40 % amplitude (4x, 1 second pulse on and 1s pulse off) for 1 minute and subsequently centrifuged at 15,000 g for 60 minutes at 4 ℃ to clear the lysate. The His-SPN fusion protein was purified with Protino Ni-Ted resin (Macherey-Nagel, Düren, Germany) using the elution buffer: 50 mM Tris, pH 7.2, 300 mM NaCl, 250 mM imidazole, 1 mM DTT, 0.1 % CHAPS, +cOMPLETE inhibitor tablet. The protein was further purified by size exclusion chromatography (SEC) with a HiLoad 26/60 Superdex 200 pg column (GE Healthcare, Chicago, IL) in SEC buffer (50 mM Tris, pH 7.2, 300 mM NaCl, 1 mM DTT, 0.1 % CHAPS) using an ÄKTA pure chromatography system (GE Healthcare). The protein was concentrated with Amicon ultracentrifugal 3K filter device (Millipore, Sigma, Burlington, MA). The homodispersity of the proteins was verified with sodium dodecyl sulphate polyacrylamide gel electrophoresis (SDS-PAGE).

#### SPN-ARR domains of SHANK3

The N-terminal SPN-ARR fragment of the rat SHANK3 (residues 1-348) cloned into pET-SUMO vector (Champion™ pET SUMO Protein Expression System, Invitrogen) was expressed and purified as described previously (2, 3).

### Atomistic simulation models and methods

#### SHANK3 with a lipid bilayer (Systems 1-3)

System S1 was comprised of SHANK3 SPN-ARR with a pure 1- palmitoyl-2-oleoyl-sn-glycero-3-phosphocholine (POPC) lipid bilayer. System S2 with SHANK3 SPN-ARR entails a symmetric three-component bilayer, containing 65 mol% POPC, 30 mol% cholesterol, and 5 mol% phosphatidylinositol 4,5-bisphosphate (PIP2). The SPN-ARR domain was initially placed ca. 2 nm away from the bilayer surface, with the SPN-ARR linker and the residues 105-115 of the ARR domain facing the bilayer. System S3 includes an isolated SPN domain (residues 2–93) together with the above-described three-component (POPC/cholesterol/PIP2) bilayer. The SPN domain was placed initially ca. 2 nm from the membrane surface.

These constructs were based on the 5G4X structure (2). Together these systems were used to probe spontaneous membrane binding capabilities of SHANK3. That is, the protein complex was initially placed in a random orientation such that the protein was allowed to bind the membrane without any bias, and these processes were simulated through four independent repeats (Extended Data Table 1). Hence, we refer to these systems as “Spontaneous” (see Extended Data Figure 3).

#### KRAS with a lipid bilayer (System S4)

System S4 entailed KRAS in an initially soluble state and a POPC/cholesterol/PIP2 lipid bilayer. The protein coordinates were extracted from the PDB id 6PTW structure (4). The resulting spontaneously formed KRAS-membrane complexes were analysed and compared to the known orientations in System S5 (see below). As in the previous case, System 4 was also studied through spontaneous binding.

#### SHANK3 and KRAS with a lipid bilayer (System S5)

<u>S</u>ystem S5 included a KRAS-SHANK3 SPN-ARR complex and a POPC/cholesterol/PIP2 lipid bilayer. The KRAS-SHANK3 complex was obtained by aligning the structures of SHANK3 SPN (2, 5) to the RAF RBD coordinates extracted from the PDB id 6PTW (4). The resulting KRAS- SHANK3 model was then equilibrated for 100 ns with 5 kJ/mol restraints on the backbone atoms. In this model, the (initial) protein-lipid configuration was extracted from the RAF RBD structure. Hence, we refer to these systems as “Model” (see Extended Data Figure 3).

Simulations were initiated using the CHARMM-GUI portal (6, 7). Interactions between the atoms were described using the all-atom CHARMM36m force field (8). Water molecules were described using the TIP3P water model (9). Potassium and chloride ions were added to neutralise the charge of the systems and to reach the physiological saline concentration (150 mM).

### Simulation parameters

To run the simulations, we used the GROMACS simulation package version 2020 (10). Initiation of the systems followed the general CHARMM-GUI protocol: the simulation systems were first energy-minimised and then equilibrated with position restraints acting on the solute atoms (8). We used the leap-frog integrator with a timestep of 2 fs to propagate the simulations (11). Periodic boundary conditions were applied in all three dimensions, atomic neighbours were tracked with the Verlet lists, and bonds were constrained with the LINCS algorithm (12). Lennard-Jones interactions were cut off at 1.2 nm, while electrostatic interactions were calculated using the smooth particle mesh Ewald (PME) algorithm (13). The pressure of the system was coupled semi-isotropically using the Parrinello-Rahman barostat with a time constant of 5 ps (13). Protein, membrane, and solvent atoms were coupled separately with a time constraint of 1 ps. Simulation trajectories were saved every 100 ps. Random initial velocities were assigned for the atoms from the Boltzmann distribution at the beginning of each simulation. For the remaining parameters, we refer to the GROMACS 2020.2 defaults (10). Production simulations are listed in Extended Data Table 1. The total simulation time of the atomistic simulations was >24 microseconds. In every system simulated, the first 100 ns were used for equilibration and were discarded from analysis. The analysis was performed for the remaining part of trajectories and over all four independent repeats/replicas (Extended Data Table 1). The error analysis, resulting in standard errors, was based on these data.

### Production of recombinant anti-SHANK3 SPN nanobodies

#### Nanobody generation

Nanobodies (single domain antibodies) against SHANK3 SPN were produced by Hybrigenics Services SAS (Evry, France; www.hybribody.com) by three rounds of Phage Display selection of their naïve VHH-library against recombinant biotinylated GST-SHANK3 SPN protein as briefly described below.

Prior to the Phage Display selection, SPN-Biotin and a non-related protein, GST-HIS-MBP-FLAG-Biotin, were bound to Streptavidin Magnetic Beads (Dynabeads® M-280 Streptavidin, Life Technologies) at a final final concentration of 50 nM (1st round) and 10 nM (2nd and 3rd rounds).

For phage display selection, unspecific binders were first removed from the hs2dAb Phage Display library by incubation with the GST-HIS-MBP-FLAG-Biotin beads. Then, the unbound VHHs expressed as an *E. coli* supernatant were incubated with the SPN-Biotin beads and a total of three rounds of Phage Display were performed. The depletion step was repeated before each round of Phage Display to remove non-specific VHHs. At the end of the 3rd round of Phage Display, *E. coli* clones were analysed by Hybrigenics’ non-adsorbed phage ELISA, which allows for the proper folding of the native SPN protein, in 384-well plates with HRP-conjugated anti- M13 antibody (GE Healthcare) and a colorimetric substrate (TMB, TetraMethylBenzidine, Thermo Fischer). VHH clones with a significant ELISA signal in the presence of SPN-Biotin and a very low signal in the presence of GST-HIS-MBP-FLAG-Biotin were considered as specific SPN binders and selected for sequence analysis. Sequencing revealed that all binders represented one of two VHH variants (hereafter called nanobodies A01 and E01). These were provided by Hybrigenics in the bacterial expression vector pHEN2 (C-terminal 6xHis and 3 Myc tags) for use in ELISA assays and in *in vitro* pulldowns, and in an mCherry mammalian expression vector (tag on C terminus) for use in cell-based assays. A single-chain variable fragment (scFv) against an unrelated protein (SorLA) was used as a negative control in ELISA assays.

#### Recombinant nanobody production

Nanobodies in the pHEN2 vector were produced as recombinant proteins in BL21 bacteria using IPTG induction and purified according to Hybrigenics’ protocol. Briefly, transformed bacteria were cultured overnight in TB buffer (tryptone 12g/L, yeast extract 24g/L, 17 mM KH2PO4, 72 mM K2HPO4, 0.4% glycerol), 1% glucose, Ampicillin 100μg/mL) at 37℃. 2 ml of this starter culture were then reincubated at 37°C until optical density at 600 nm was between 0.6 and 0.8, induced with IPTG (0.5 mM) overnight at 28°C, pelleted and freeze-thawed in liquid nitrogen. The pellet was resuspended in sonication buffer (50 mM NaPO4, pH 8, 300 mM NaCl, bacterial protease inhibitors, 1 mM PMSF, 1mg/ml lysozyme), sonicated on ice (3 times for 1 second) and centrifuged to remove debris. Lysates were then incubated with prewashed talon metal affinity resin (BD) at 4°C for 30 minutes with shaking. Resin was then spun, flow-through removed, and washed with sonication buffer. Nanobodies were then eluted in sonication buffer + 250 mM imidazole.

### Functional ELISA assay for nanobody testing

Nunc maxisorp 96-well plates were coated with 5 ug/ml purified recombinant His-SHANK3 SPN protein in TBS, 100 l/well, overnight at 4 ℃. Wells coated with BSA alone were included as a background binding control. The coating solution was removed, the wells were blocked with 100 l/well 5% BSA in TBS-0.1% Tween20 (TBSt) for 1 h at room temperature. The His-SPN coated wells were preincubating with nanobodies for 15 minutes in room temperature prior to addition of GST-tagged purified recombinant KRAS protein loaded with non-hydrolysable GTP analogue. An irrelevant anti-SorLA single-chain antibody (sc-Fv) was added as a negative control. These were incubated for 1 h at room temperature in TBSt+1 mM DTT+2 mM MgCl2. Wells were washed 3 times with TBSt+2 mM MgCl2. To detect GST, DELFIA® Eu-N1 Anti-GST antibody (Perkin Elmer catalogue number AD0250) diluted at 1:000 in TBSt+1 mM DTT+2 mM MgCl2 was added l/well and incubated 1h at room temperature. The wells were washed 3 times with Tecan plate washer with PBS, 100 ul of DELFIA® Enhancement Solution (Perkin Elmer catalog number 124-105) was added to the wells and europium signal was measured with a time resolved fluorescence plate reader (PerkinElmer’s VICTOR X5 multilabel plate reader).

### Pull-down interaction assay

The ability of anti-SHANK3 SPN nanobodies (A01 and E01) to disrupt the interaction between SHANK3 SPN and KRASG12V was tested with a pull-down. 5 ug of His-SHANK3 SPN protein was bound to 20 ul of Macherey Nagel Protino Ni-Ted resin beads in TBS+1 mM DTT+2 mM MgCl2 for 1h under rotation at 4 ℃. 20 ug of nanobodies A01 or E01 or BSA as control were incubated with this for 30 min under rotation under rotation at 4 ℃. 5 ug/ml GST-KRAS-GTP or GST alone was added and incubated under rotation at 4 ℃ for 1h. Beads were washed 3 times with 500 ul TBS+1 mM DTT+2 mM MgCl2, eluted into 20 ul 4x SDS sample buffer (200 mM Tris-HCl pH 6.8, 8% SDS, 40% glycerol, 4% β-mercaptoethanol, 50 mM EDTA, 0.08 % bromophenol blue) with heating at 90 ℃ for 5 min and separated on an SDS-PAGE gel. The proteins were transferred to a filter and western blotted with anti-GST antibody.

